# Mitotic CDK promotes replisome disassembly, fork breakage, and complex DNA rearrangements

**DOI:** 10.1101/428433

**Authors:** Lin Deng, R. Alex. Wu, Olga V. Kochenova, David Pellman, Johannes C. Walter

## Abstract

DNA replication errors generate complex chromosomal rearrangements and thereby contribute to tumorigenesis and other human diseases. Although the events that trigger these errors are not well understood, one candidate is mitotic entry before the completion of DNA replication. To address the impact of mitosis on DNA replication, we employed *Xenopus* egg extracts. When mitotic CDK (Cyclin B1-CDK1) is used to drive these extracts into mitosis, the E3 ubiquitin ligase TRAIP promotes ubiquitylation of the replicative CMG (<u>C</u>DC45/<u>M</u>CM2–7/<u>G</u>INS) helicase at stalled forks and at forks that have completed DNA synthesis. In both cases, ubiquitylation is followed by CMG extraction from chromatin by the CDC48/p97 ATPase. At stalled forks, CMG removal results in fork breakage and complex end joining events involving deletions and template-switching. Our results identify TRAIP-dependent replisome disassembly as a novel trigger of replication fork collapse and propose it underlies complex DNA rearrangements in mitosis.

**HIGHLIGHTS:**

1. TRAIP-dependent MCM7 ubiquitylation removes all CMGs from chromatin in mitosis
2. CMG unloading from stalled forks causes replication fork breakage
3. Replication fork breakage in mitosis causes complex rearrangements
4. New model of replication fork collapse

## INTRODUCTION

Genome evolution occurs through the gradual accrual of genetic changes or in a saltatory manner, with bursts of chromosomal alterations originating from single catastrophic events (Holland and Cleveland, 2012; Leibowitz et al., 2015; Liu et al., 2011; Stephens et al., 2011). Many chromosomal alterations can be traced to DNA breaks that arise during DNA replication (Hills and Diffley, 2014; Mankouri et al., 2013; Techer et al., 2017). However, there is an ongoing debate about when and how replication fork breakage is triggered (Toledo et al., 2017).

In normal cells, multiple cell cycle regulatory controls and error correction mechanisms prevent DNA replication errors (Hills and Diffley, 2014). Cells prepare for DNA replication in the G1 phase of the cell cycle, when pairs of MCM2–7 ATPases are recruited to each origin (“licensing”). In S phase, cyclin-dependent kinase (CDK) promotes the association of CDC45 and GINS with MCM2–7, leading to formation of the replicative CMG helicase complex (<u>C</u>DC45-<u>M</u>CM2–7-<u>G</u>INS) (“initiation”). CMG unwinding of the origin nucleates the assembly of two DNA replication forks that travel away from the origin, copying DNA as they go (“elongation”). When converging forks from adjacent origins meet, the replisome is disassembled (“termination”). Replisome disassembly in S phase requires the E3 ubiquitin ligase CRL2^Lrr1^, which ubiquitylates the MCM7 subunit of CMG, leading to CMG’s extraction from chromatin by the p97 ATPase (Dewar et al., 2017; Sonneville et al., 2017). In the absence of CRL2^Lrr1^, CMGs persist on chromatin until mitosis, but are then removed by a secondary, p97-dependent pathway that is controlled by an unknown E3 ubiquitin ligase (Sonneville et al., 2017). Re-replication is inhibited because *de novo* licensing of origins is suppressed in the S and G2 phases of the cell cycle. In summary, faithful DNA replication requires the seamless integration of replication licensing, initiation, elongation, and termination. Errors in the process are detected by the DNA damage response, which activates repair mechanisms and prevents entry into mitosis in the setting of incomplete or abnormal replication.

DNA replication forks become stressed in a variety of circumstances, including the activation of oncogenes, the presence of replication-blocking DNA lesions, and nucleotide starvation (Cortez, 2015; Hills and Diffley, 2014; Saldivar et al., 2017). Replication stress, especially when combined with inhibition of checkpoint kinases, can cause replication fork “collapse”, which is defined as an irreversible state from which replication cannot be restarted (Cortez, 2015; Hills and Diffley, 2014; Pasero and Vindigni, 2017; Saldivar et al., 2017). Numerous experiments in different eukaryotic organisms indicated that fork collapse involves replisome disassembly (Cortez, 2015). However, these studies did not determine which component(s), when lost, trigger collapse, and they did not establish a causal relationship between replisome disassembly and collapse. More recent experiments suggest that fork collapse may not involve replisome disassembly (Cortez, 2015; De Piccoli et al., 2012). In some cases, fork collapse is accompanied by breakage of DNA at the fork, but the relationship between these two processes is unclear. In summary, there is little agreement on the basic processes that underlie the irreversible inactivation of DNA replication forks.

In vertebrates, the checkpoint kinase ATR protects forks from collapse, but the underlying mechanism has also been vigorously debated (Toledo et al., 2017). A widespread view is that ATR promotes the phosphorylation of unspecified proteins at forks to prevent their collapse (Cortez, 2015). Another model is that the key function of ATR is to restrain cell cycle progression. One version of this idea is that in the absence of ATR, excessive origin firing leads to exhaustion of the nuclear pool of RPA, followed by fork breakage and replisome collapse (Toledo et al., 2013). Alternatively, ATR might prevent fork collapse by restraining mitotic entry, which would delay the activation of mitotic kinases such as PLK1 (Ragland et al., 2013). In agreement with the latter model, genetic studies suggest that, in mammals, restraining cell cycle progression is the essential function of ATR (Ruiz et al., 2016). Mitotic kinases can induce fork breakage by promoting the assembly of a MUS81-containing nuclease complex (Duda et al., 2016) or by triggering nuclear envelope breakdown, which grants the normally cytoplasmic GEN1 nuclease access to replication forks (West and Chan, 2018). Thus, there is no consensus on whether ATR prevents fork collapse primarily by local control at the fork or via restraint of cell cycle progression.

Replication fork breakage is sometimes beneficial. A prominent example involves commons fragile sites (CFS), genomic loci that replicate late in S phase and are difficult to replicate because they contain few origins of replication and large genes with long transcripts (Glover et al., 2017). Common fragile site “expression,” the appearance of cytologically visible breaks and gaps, is promoted by low doses of aphidicolin because this drug delays duplication of already late-replicating loci. CFS are also among the most frequently rearranged loci in cancer genomes. In aphidicolin-treated cells, CFS co-localize with ultrafine DNA bridges that link the separating sister chromatids in anaphase (Baumann et al., 2007; Chan et al., 2007). Depletion of MUS81 exacerbates these segregation errors, inhibits CFS expression, and increases the frequency of “53BP1 bodies” (Naim et al., 2013; Ying et al., 2013), structures thought to protect damaged DNA at these sites in the ensuing interphase (Harrigan et al., 2011; Lukas et al., 2011). Collectively, the data suggest that when cells enter mitosis before completion of DNA replication at CFS, MUS81 promotes breakage of stalled replication forks. This enables chromosome segregation, but comes with the risk of forming deletions and other rearrangements. These findings indicate that CFS expression is beneficial. However, no model has emerged that explains how CFS expression avoids deleterious outcomes such as the generation of acentric or iso-chromosomes.

Although breakage of a few stressed forks can be beneficial, concurrent breakage of many forks is deleterious as it leads to catastrophic chromosomal rearrangements. Several lines of evidence also implicate mitotic entry as one potential cause of extensive fork breakage. Cell fusion experiments (Johnson and Rao, 1970) and experiments on cells with micronuclei (Kato and Sandberg, 1968) showed that S phase chromosomes undergo “pulverization” upon exposure to mitotic cytoplasm. Although there was early disagreement about whether chromosome pulverization reflects discontinuous condensation or actual DNA breakage (Rao et al., 1982), recent work indicates that fragmentation does occur. First, premature mitotic entry triggered by inhibition of the WEE1 kinase causes extensive fork breakage in a manner that depends upon SLX4 and MUS81 (Dominguez-Kelly et al., 2011; Duda et al., 2016). Second, chromothripsis, a mutational process involving extensive chromosome fragmentation and rearrangement, may involve entry into mitosis of micronuclei undergoing DNA replication (Crasta et al., 2012; Leibowitz et al., 2015). Extensive fork breakage during mitosis is problematic as both homologous recombination and classical non-homologous end joining are inactive during mitosis (Hustedt and Durocher, 2016). In summary, it has become apparent that genome instability in a variety of contexts is linked to replication fork breakage during mitosis. However, the general question of how mitosis impacts the normal program of DNA replication remains poorly understood.

Here, we used *Xenopus* egg extracts to explore the relationship between DNA replication and mitosis. We find that when extracts containing stressed replication forks are driven into mitosis with Cyclin B1-CDK1, the CMG helicase is ubiquitylated on its MCM7 subunit and subsequently extracted from chromatin by the CDC48/p97 ATPase. We show that the E3 ubiquitin ligase TRAIP is critical for this pathway. TRAIP-dependent CMG unloading leads to fork breakage, followed by end joining events that likely involve DNA polymerase θ (Polθ). Importantly, in the absence of CRL2^Lrr1^, TRAIP also promotes the removal of CMGs from replisomes that have undergone replication termination, indicating that TRAIP clears the chromosomes of all CMGs in mitosis. Together, our results identify CMG loss from the fork as a new mechanism of replication fork collapse and ultimately fork breakage. We propose that breakage of a few converging forks that have failed to complete DNA synthesis in mitosis helps to maintain chromosome integrity whereas breakage of many forks leads to catastrophic rearrangements.

## RESULTS

### Mitotic CDK triggers aberrant processing of stressed DNA replication forks

To examine the effect of mitotic CDK on DNA replication, we employed *Xenopus* egg extracts, which can be permanently arrested in states that mimic S phase or mitosis while also allowing careful monitoring of DNA replication forks. To this end, plasmid DNA was incubated in a high-speed supernatant (HSS) of *Xenopus* egg extract. HSS promotes the assembly onto DNA of pre-replication complexes (pre-RCs) containing double hexamers of the MCM2–7 ATPase (Figure 1A). The subsequent addition of a nucleoplasmic extract (NPE) leads to the association of CDC45 and GINS with each MCM2–7 hexamer to form two active CMG DNA helicases, which unwind DNA at the fork, promoting a single, complete round of DNA replication (Figure 1B, lanes 1–6) (Walter et al., 1998). Because the mitotic CDK Cyclin B1-CDK1 (B1-CDK1) inhibits licensing in egg extract (Hendrickson et al., 1996; Prokhorova et al., 2003), we added it after pre-RC formation (Figure 1A) at a concentration that induces chromosome condensation (Figures S1A-S1C) and condensin recruitment (Figures S1D and S1E). As we showed previously (Prokhorova et al., 2003), B1-CDK1 increased the rate of DNA replication in NPE (Figure 1B, compare lanes 1–6 and 13–18). This acceleration was due in part to increased origin firing, as shown by enhanced CMG loading (Figure S1F). However, in the absence of other perturbations, all replication products were open circular or supercoiled species (Figure 1B, lanes 13–18), indicating that B1-CDK1-induced chromatin condensation does not cause aberrant DNA replication.

**Figure 1.**
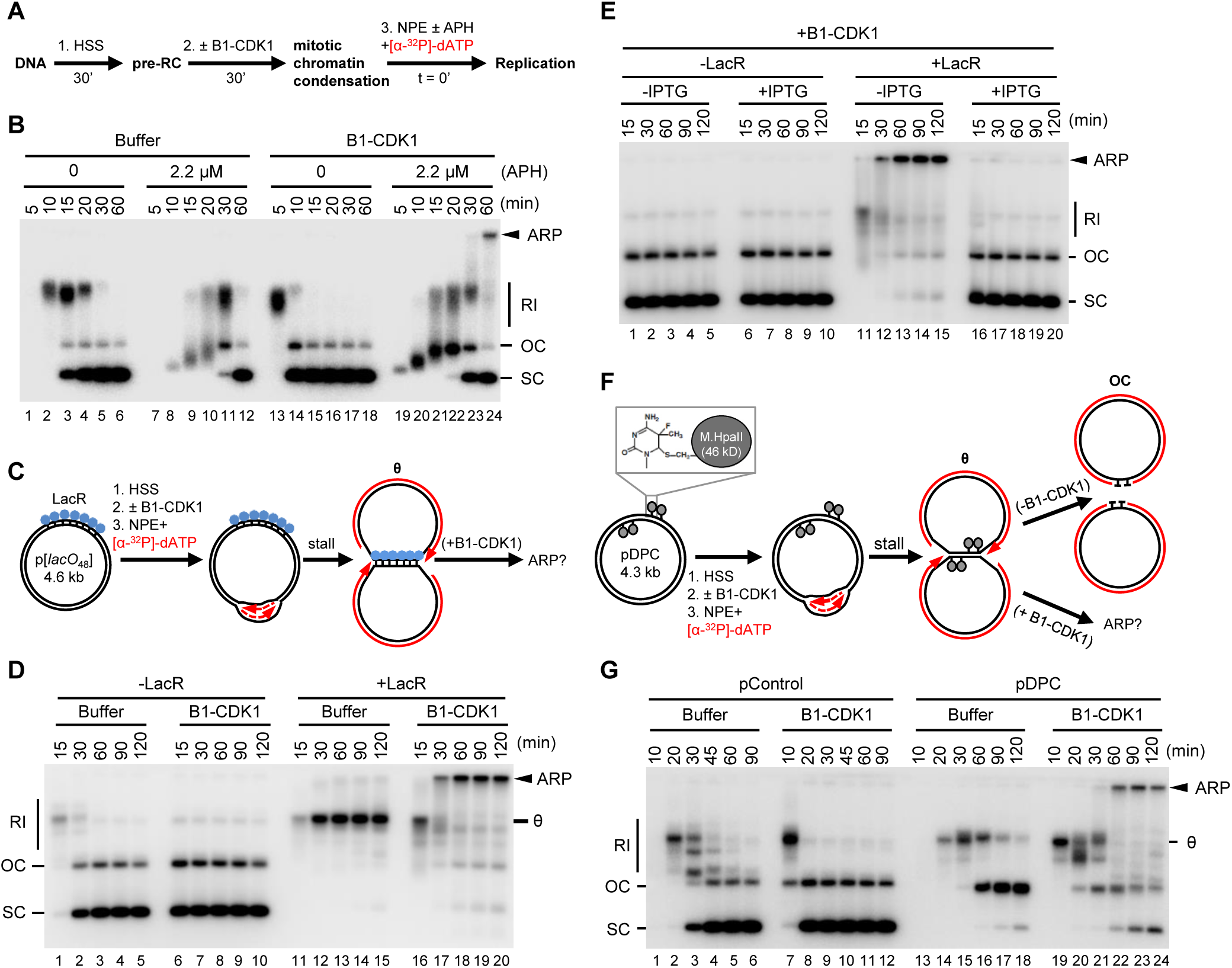
Mitotic CDK triggers aberrant processing of stalled DNA replication forks in *Xenopus* egg extracts. **(A)** Schematic of experimental approach to test effect of B1-CDK1 on DNA replication. APH, DNA polymerase inhibitor aphidicolin. **(B)** A 3 kb pBlueScript plasmid was replicated according to (A) and products were separated on a native agarose gel followed by autoradiography. Unless stated otherwise, the ‘0 minute’ time point refers to NPE addition. **(C)** Schematic of DNA replication for LacR-bound p[*lacO***_48_**] plasmid. **(D)** p[*lacO***_48_**] was replicated according to (C) under the indicated conditions. **(E)** p[*lacO***_48_**] was replicated according to (C) in the presence of LacR and/or IPTG (10 mM, 15 min incubation in NPE before mixing with “licensing” mixture), as indicated. **(F)** Schematic of replication for pDPC, containing four 46 kDa M.HpaII DNA methyltransferases at the indicated positions. Products formed in the presence and absence of B1-CDK1 are indicated. **(G)** pControl or pDPC was replicated according to (F) using the indicated conditions. From (A) to (G), B1-CDK1 was added to “licensing” mixture at a concentration of 50 ng/μL and its final concentration in the overall reaction is 16.7 ng/μL. RI, replication intermediate; OC, open circle; SC: supercoil; θ, theta structure; ARP, aberrant replication product. See also **Figure S1**.

Given the evidence that stressed DNA replication forks undergo breakage during mitosis (e.g. at common fragile sites, see introduction), we added a low concentration of the replicative DNA polymerase inhibitor aphidicolin (APH; 2.2 μM) to slow fork progression (Figure 1B, lanes 7–12). Intriguingly, the combination of B1-CDK1 and APH (Figure 1B, lanes 19–24) led to the appearance of a new replication product that migrated at the very top of the gel. This aberrant replication product (ARP) comprised ~6% of total replication for a 3 kb plasmid and up to 30% for a 9 kb plasmid (data not shown), presumably because the larger plasmid hosts more replication forks. The ARPs were not resolved by Topoisomerase I or Topoisomerase II treatment, indicating they are not plasmid topoisomers (data not shown). Thus, in the presence of replication stress, mitotic CDK induces aberrant DNA replication.

To mimic incomplete DNA replication in mitosis, as occurs at common fragile sites, we stalled replication forks on either side of defined replication fork barriers before addition of B1-CDK1. First, we replicated a plasmid containing an array of 48 *lacO* sites (p[*lacO***_48_**]) bound by the *lac* repressor (LacR) (Figure 1C). As expected (Dewar et al., 2015), replication forks stalled at the outer edges of the LacR array, generating a “theta” (θ) structure (Figures 1C and 1D, lanes 11–15). Strikingly, in the presence of B1-CDK1, the theta molecules disappeared and ARPs accumulated (Figure 1D, lanes 16–20). ARPs were not generated when LacR-mediated fork stalling was prevented with IPTG (Figure 1E), or in the presence of the CDK1 inhibitor (CDK1-i) RO-3306 (Figure S1G). Furthermore, the S-phase CDK, Cyclin E-CDK2, did not induce ARPs (data not shown). Second, we induced replication fork stalling with covalent DNA-protein crosslinks (DPCs). We replicated a plasmid substrate (pDPC), which contains two site-specific DPCs on each leading strand template (Figure 1F). As expected (Duxin et al., 2014), in the absence of B1-CDK1, replication of pDPC first yielded theta structures when forks transiently paused at the DPC. Plasmids then resolved into open circular (OC) species that persisted due to slow translesion synthesis past the peptide adduct generated by DPC proteolysis (Figure 1F, upper arrow and Figure 1G, lanes 13–18). In the presence of B1-CDK1, we again observed a substantial accumulation of ARPs (Figure 1G, lanes 19–24). In summary, mitotic CDK caused aberrant processing of replication forks stalled by aphidicolin, non-covalent nucleoprotein complexes, and DPCs.

### Mitotic processing of stalled replication forks leads to complex chromosomal rearrangements

To determine the structure of mitotic ARPs, we replicated the 4.6 kb LacR plasmid in the presence and absence of B1-CDK1 and digested the replication products with AlwNI and AflII, which cuts the plasmid into a 1.9 kb fragment and a 2.7 kb fragment encompassing the *lacO* repeats (Figure 2A). In the absence of B1-CDK1, fully replicated 1.9 kb fragments quickly accumulated, whereas the rest of the plasmid migrated as a double-Y structure that gradually increased in size due to slow progression of forks through the LacR array (Figure 2B, middle panel, lanes 1–7 and Figure 2C; see also (Dewar et al., 2015) Figure S4). In the presence of B1-CDK1, the 1.9 kb fragment again accumulated quickly and persisted, demonstrating that this *lacO*- free region was replicated efficiently (Figure 2B, middle panel, lanes 8–14). However, the double-Y structure containing the *lacO* array rapidly disappeared. Thus, in the presence of B1-CDK1, DNA processing occurs specifically on molecules containing stalled forks.

**Figure 2.**
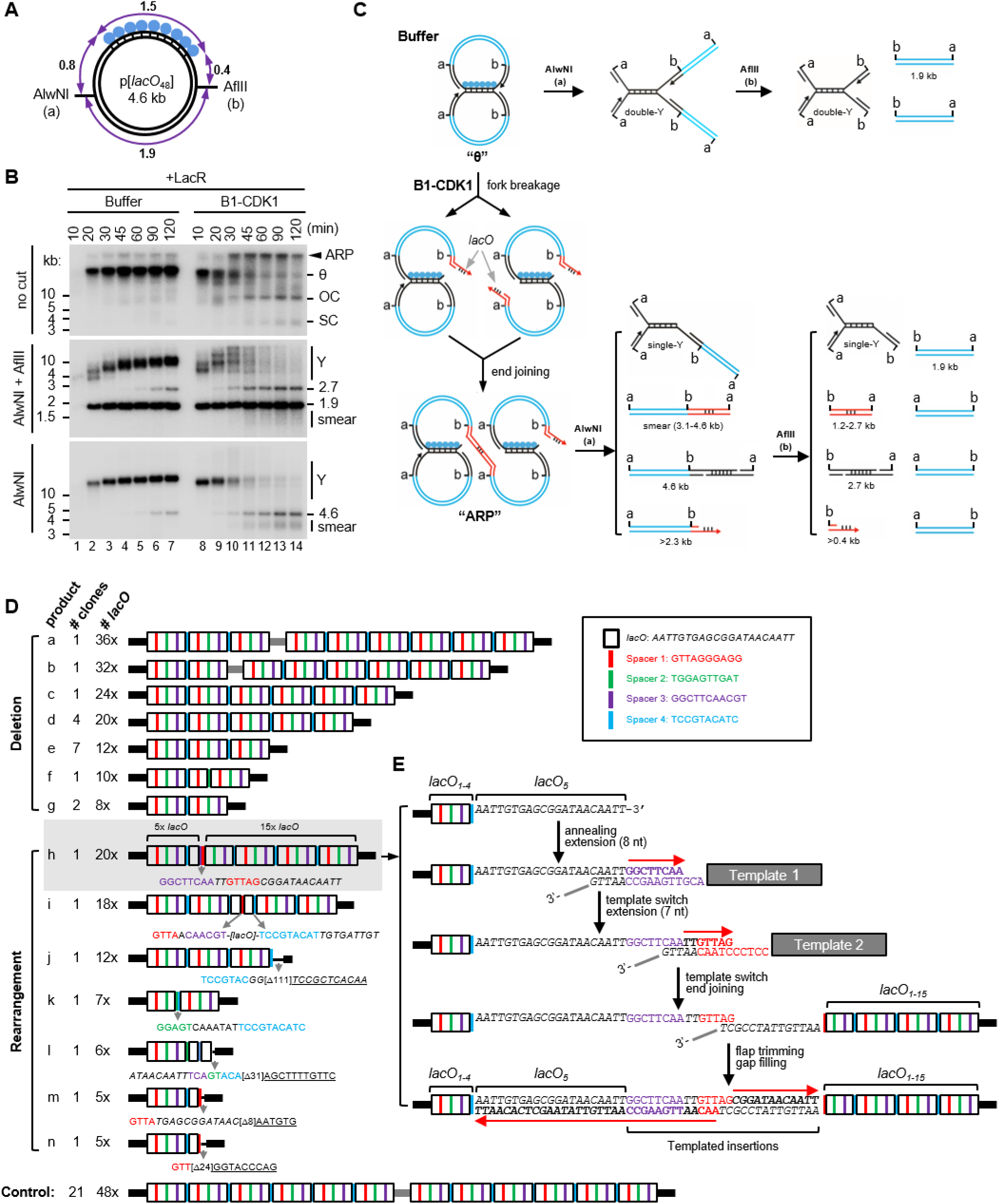
Mitotic processing of stalled replication forks leads to complex DNA rearrangements. **(A)** Structure of the 4.6 kb p[*lacO***_48_**] plasmid. Numbers mark the length of the indicated DNA segments in kilo-basepairs (kb). **(B)** p[*lacO***_48_**] was replicated in the presence of Buffer or B1-CDK1. At the indicated time points, replication products were isolated and digested with AlwNI and AflII, or AlwNI, as indicated. Numbers label the size of linear fragments in kb; Y, double-Y or single-Y structure (see panel C). **(C)** Model explaining the restriction products observed in (B). Although the model favor the fork breakage on the leading strand, the possibility of fork breakage on the lagging strand has not been excluded. A more detailed model is in Figure S2A. **(D)** The smear of ~3–4 kb mitotic DNA replication products generated after AlwNI digestion in (B) was self-ligated, cloned and sequenced. The controls are replication products of the same plasmid from a mitotic reaction lacking LacR. The *lacO* repeats, shown as white boxes, are separated by four unique spacers shown in different colors. Inset, DNA sequences of the *lacO* repeat and four spacers. The detailed structure of the entire *lacO* array is shown in Figure S2C. **(E)** A model for the generation of product h in (D) from multiple template-switching events. See also **Figure S2**.

When the replication products were digested only with AlwNI, we observed B1-CDK1-dependent disappearance of the now larger double-Y structure (Figure 2B, bottom panel, lanes 8–14). In addition, we detected a new series of species migrating between ~3 and ~4 kb (Figure 2B, bottom panel; smear). We hypothesized that when replication forks enter the array and slow down or stall, B1-CDK1 promotes their collapse and breakage. The resulting double-strand breaks (DSBs) subsequently undergo joining with DSBs from broken forks on other plasmids, generating ARPs (Figures 2C and S2A). If replication forks collapse at the outer edges of the array, the size of the end joining product after AlwNI digestion is close to 3.1 kb because most of the 1.5 kb *lacO* array is lost; collapse further into to the array generates larger products, accounting for the 3–4 kb range of products observed (Figure S2B). To test this hypothesis, the 3–4 kb species were cloned and sequenced using primers immediately flanking the *lacO* array (Figure S2C). In contrast to control clones (generated from replication in the absence of LacR), all of which contained 48 *lacO* repeats, the 24 clones from the 3–4 kb smear contained fewer than 48 *lacO* repeats (Figure 2D, products a-n). This result confirms that replication forks collapsed within the *lacO* array and then underwent end joining with loss of *lacO* repeats. Seventeen of these products (a-g) involved simple deletions of the *lacO* repeats. Repeats were mostly lost in blocks of four *lacO* sites, the repeating unit within the *lacO* array that also contains four unique spacer sequences. This suggests that the deletions might occur via single strand annealing (SSA) (Bhargava et al., 2016), which generates deletions between homologous sequences of this length. The remaining 7 clones had complex rearrangements, including insertions that likely arose from replication template-switching events (Figure 2D; product h-n). For example, product h appears to have arisen from fork collapse at the 5^th^ repeat, followed by two successive microhomology-mediated strand invasion and copying events, followed by joining to a second fork that broke at the 15^th^ repeat (Figure 2E). Together, the sequencing data strongly suggest that stressed replication forks collapse in the presence of B1-CDK1, generating DSBs that subsequently undergo end joining (Figures 2C and S3A), sometimes after repeated template-switching (Figure 2E).

### Immunodepletion of DNA Polθ reduces mitotic ARPs

We next addressed the mechanism of end joining after mitotic CDK-induced fork collapse. As expected (Hustedt and Durocher, 2016; Peterson et al., 2011), RAD51, which is essential for homologous recombination (HR), did not bind chromatin in the presence of B1-CDK1 (Figure S3A). Accordingly, immunodepletion of RAD51 from egg extracts had no effect on B1-CDK1-induced ARP formation (Figures S3B and S3C), nor did inhibition of RAD51 with a BRC peptide derived from BRCA2 (Figure S3D) (Long et al., 2011). Further, classical non-homologous end joining (NHEJ), which is also normally inhibited during mitosis (Hustedt and Durocher, 2016), was not required for ARP formation (Figure S3E). The structures of the mitotic ARPs (Figures 2C-2E) suggested that MMEJ (microhomology-mediated end joining, also called alternative end joining) and/or SSA might be responsible for mitotic DSB repair. Indeed, immunodepletion of DNA polymerase Polθ (Figure 3A), a major mediator of MMEJ known to make errors due to replicative template-switching (Wyatt et al., 2016), decreased ARPs during replication of LacR plasmid (Figures 3B and S3F) and pDPC (Figures 3C and S3G). Although we have so far not rescued this effect with purified Polθ protein, the involvement of Polθ is consistent with the nature of the end joining products shown in Figures 2D-2E. Thus, in mitotic extracts where HR and NHEJ are inactive, MMEJ appears to become a major pathway that mediates joining of DNA ends after fork breakage.

**Figure 3.**
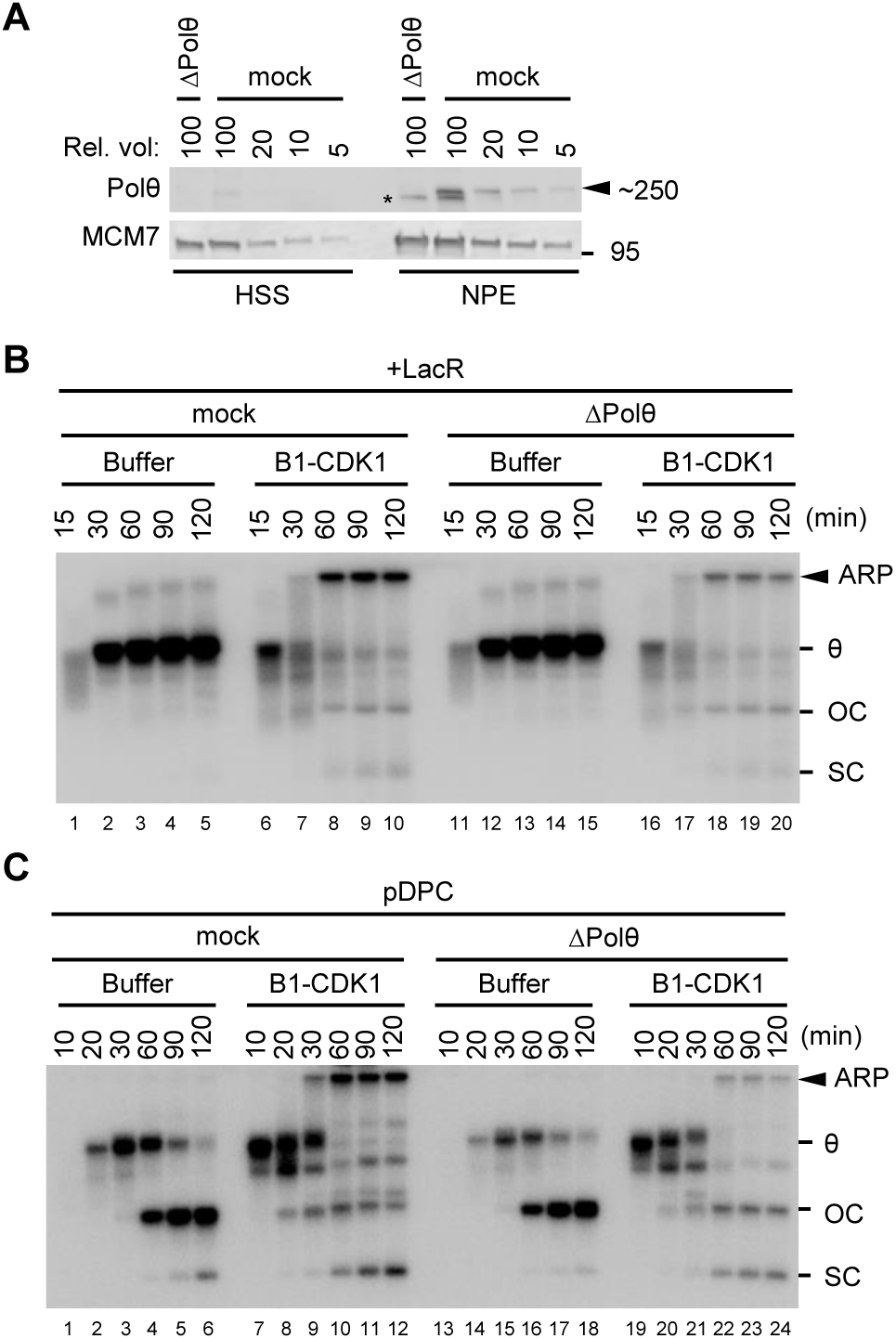
Depletion of DNA polymerase θ disrupts the generation of aberrant replication product in mitosis. **(A)** Mock-depleted and Polθ-depleted *Xenopus* egg extracts were blotted for Polθ and MCM7, alongside a serial dilution of mock-depleted extracts. Asterisk, background band. **(B)** LacR-bound p[*lacO***_48_**] was replicated in mock-depleted or Polθ-depleted extracts with or without B1-CDK1 treatment. Overall DNA replication and ARP were quantified in Figure S3F. **(C)** pDPC was replicated in mock-depleted or Polθ-depleted egg extracts with or without B1-CDK1 treatment. Overall DNA replication and ARP were quantified in Figure S3G. In (B) and (C), OC, open circle; SC, supercoil; θ, theta structure; ARP, aberrant replication product. See also **Figure S3**.

### Chromatin condensation does not cause fork breakage

We next sought to address how mitotic CDK causes fork instability. Chromatin condensation, a central event in mitosis, has long been proposed to cause DNA damage in under-replicated regions (El Achkar et al., 2005; Lukas et al., 2011). We therefore examined the role of chromatin condensation on mitotic fork collapse. Although immunodepletion of the condensin subunit SMC2 inhibited B1-CDK1-induced chromosome condensation (Figures S4A-B), it did not affect the formation of ARPs (Figures 4A and S4C). This finding is consistent with our results that condensin recruitment did not induce DNA damage in the absence of replication stress (Figures 1B, 1D, 1G and S1C-S1E). These data indicate that chromatin condensation, *per se*, is neither necessary nor sufficient for fork instability in mitotic egg extracts.

**Figure 4.**
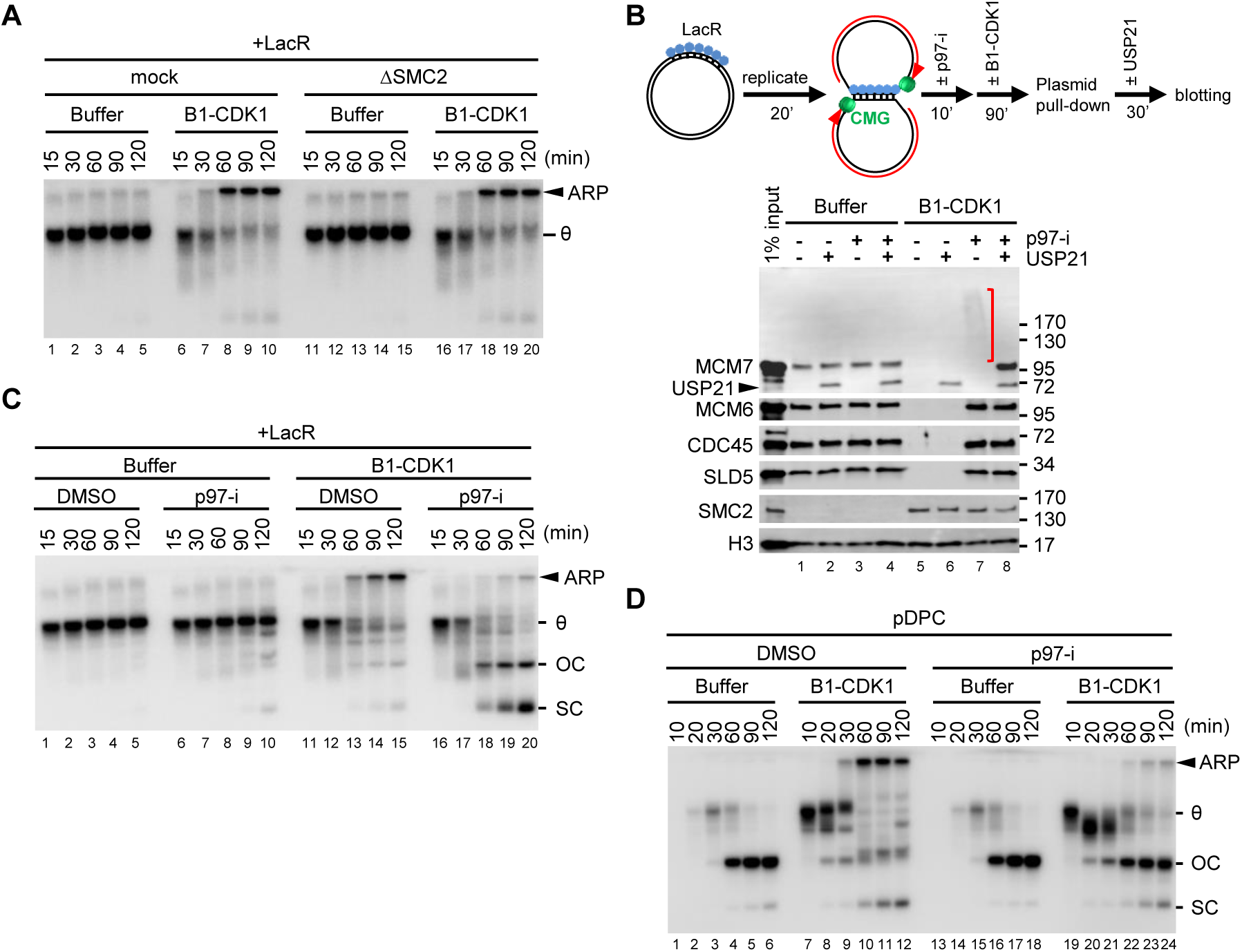
Mitotic replication fork collapse requires p97-dependent CMG unloading. **(A)** LacR-bound p[*lacO***_48_**] was replicated in mock-depleted or condensin SMC2-depleted extracts with or without B1-CDK1 treatment. **(B)** LacR-bound p[*lacO***_48_**] plasmid was replicated and treated as schemed. Chromatin-bound proteins were recovered and blotted with the indicated antibodies. Red bracket, ubiquitylated MCM7. Histone H3 served as a loading control. Note that the MCM7 antibody cross-reacts with USP21. **(C)** LacR-bound p[*lacO***_48_**] was replicated in the presence or absence of p97-i and B1-CDK1, as indicated. **(D)** pDPC was replicated in the presence or absence of p97-i and B1-CDK1, as indicated. ARP, OC+SC and overall DNA replication were quantified in Figure S4E. In (A), (C) and (D), OC, open circle; SC: supercoil; θ, theta structure; ARP, aberrant replication product. See also **Figure S4**.

### CMG unloading at stalled forks initiates mitotic fork breakage

When replication forks stall on either side of a DNA inter-strand crosslink (ICL) in interphase egg extracts, CMGs are ubiquitylated and unloaded from chromatin by the CDC48/p97 ATPase (Fullbright et al., 2016; Semlow et al., 2016). The loss of CMGs from the stalled forks enables XPF-dependent ICL incision (Klein Douwel et al., 2014), which unhooks the lesion, leading to the formation of a double-stranded DNA break that is subsequently repaired via homologous recombination (Long et al., 2014). Inspired by this mechanism, we asked whether B1-CDK1-induced fork breakage at single stalled forks is caused by CMG unloading.

As shown previously (Dewar et al., 2015), CMGs that stalled at a LacR array did not dissociate from chromatin in interphase extracts (Figure 4B, lane 1). Intriguingly, in the presence of B1-CDK1, CMGs were unloaded efficiently (Figure 4B, lane 5). Addition of the p97 inhibitor NMS-873 (p97-i) prevented B1-CDK1-triggered CMG unloading and revealed a ladder of MCM7 species (Figure 4B, lane 7, red bracket) that was collapsed by USP21, a non-specific deubiquitinating enzyme (Figure 4B, lane 8). Therefore, B1-CDK1 induces MCM7 ubiquitylation and CMG unloading from single stalled forks, in the absence of replication fork convergence. Strikingly, p97-i suppressed the formation of ARPs on the LacR plasmid (Figure 4C; see below for explanation of OC and SC product formation), strongly suggesting that B1-CDK1-induced CMG unloading triggers replication fork breakage. Consistent with this interpretation, CMG unloading normally preceded replication fork breakage (Figure S4D). As seen for LacR plasmid, p97-i also prevented ARP formation on pDPC (Figures 4D and S4E). Our data demonstrate that breakage of stalled DNA replication forks in the presence of mitotic CDK requires p97 activity, likely due to its role in CMG unloading.

### TRAIP promotes CMG unloading from stalled forks in mitosis

We next sought to identify the E3 ubiquitin ligase that promotes MCM7 ubiquitylation in mitosis. Although CRL2^Lrr1^ normally acts on CMGs that encircle dsDNA after passing each other during replication termination (Dewar et al., 2017), it was possible that B1-CDK1 might target it to stalled CMGs that encircle ssDNA. However, while the Cullin inhibitor MLN-4924 (Cul-i) blocked CMG unloading during replication termination in interphase (Figure S5A, compare lanes 1 and 4), it had almost no effect on mitotic CMG unloading from stalled forks (Figure S5A, compare lanes 3 and 6), indicating the latter process does not involve CRL2^Lrr1^. Therefore, a Cullin-independent E3 ubiquitin ligase is responsible for MCM7 ubiquitylation upon premature mitotic entry.

The E3 ubiquitin ligase TRAIP counteracts replication stress to maintain genome integrity (Feng et al., 2016; Harley et al., 2016; Hoffmann et al., 2016; Soo Lee et al., 2016), and we recently found that it is bound to replication forks that have stalled at a LacR array (Dewar et al., 2017). We therefore asked whether TRAIP is responsible for CMG unloading from stalled forks in mitosis. Strikingly, immunodepletion of TRAIP from egg extract (Figure 5A) prevented B1-CDK1-induced CMG unloading at stalled forks (Figure 5B, compare lanes 2 and 6), and it eliminated the polyubiquitylation of MCM7 detected in the presence of p97-i (Figure 5B, compare lanes 4 and 8). Furthermore, TRAIP depletion abolished the formation of ARPs during replication of LacR plasmid (Figure 5C, compare lanes 7–12 and 19–24) and pDPC (Figure S5B). To determine whether the effect of TRAIP depletion was specific, we purified recombinant wild type TRAIP protein (rTRAIP^WT^) from bacteria (Wu et al., submitted). Addition of rTRAIP^WT^ to TRAIP-depleted egg extracts rescued formation of mitotic ARPs (Figure 5D, compare lanes 13–18 to 7–12; and Figures S5C-S5E). We also added back TRAIP carrying the substitution R18C, which was identified in a human patient with primordial dwarfism (Harley et al., 2016) and that severely reduces the E3 ligase activity of TRAIP (Wu et al., submitted). Unlike rTRAIP^WT^, rTRAIP^R18C^ supported only low levels of ARP formation on LacR plasmid (Figure 5D, compare lanes 19–24 to 13–18). We conclude that TRAIP is essential for replication fork collapse at stalled forks in mitosis, most likely due to a role in MCM7 ubiquitylation and p97-dependent CMG unloading.

**Figure 5.**
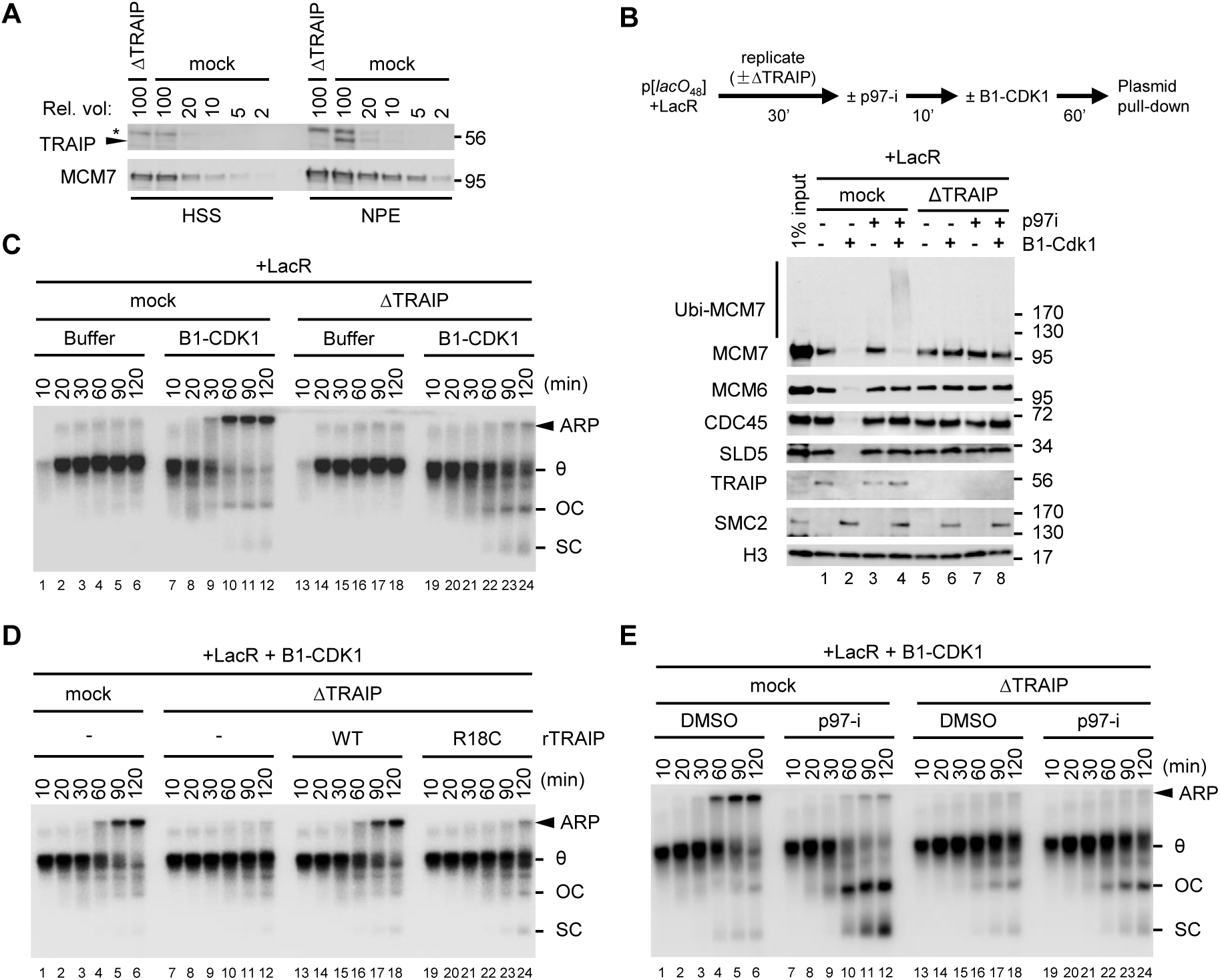
E3 ubiquitin ligase TRAIP promotes mitotic CMG unloading from a stalled replication fork. **(A)** Mock-depleted and TRAIP-depleted egg extracts were blotted for TRAIP and MCM7 alongside a serial dilution of mock-depleted extracts. **(B)** LacR-bound p[*lacO***_48_**] plasmid was replicated in mock-depleted or TRAIP-depleted egg extracts and treated as schemed. Chromatin-bound proteins were recovered and blotted with the indicated antibodies. **(C)** LacR-bound p[*lacO***_48_**] was replicated in mock-depleted or TRAIP-depleted extracts with or without B1-CDK1 treatment. **(D)** LacR-bound p[*lacO***_48_**] was replicated in mitotic mock-depleted or TRAIP-depleted egg extracts with or without recombinant rTRAIP^WT^ or rTRAIP^R18C^, as indicated. rTRAIP^WT^ and rTRAIP^R18C^ were added to NPE at a concentration of 21 ng/μL (~7-fold of endogenous TRAIP, see assessment in Figure S5C). Matched buffer was added to reactions without recombinant protein. Note that addition of rTRAIP^WT^ at endogenous level into TRAIP-depleted extracts also strongly rescued mitotic ARPs, see Figures S5D and S5E. **(E)** LacR-bound p[*lacO***_48_**] was replicated in mock-depleted or TRAIP-depleted mitotic egg extracts with DMSO or p97-i treatment. See also **Figure S5**.

To understand how TRAIP is regulated, we monitored its binding to chromatin. As we showed previously (Dewar et al., 2017), in interphase egg extract TRAIP is associated with replisomes that have stalled at a LacR array (Figure 5B, lane 1). Therefore, TRAIP is present at forks before they are exposed to B1-CDK1. Upon addition of B1-CDK1, TRAIP was lost from the chromatin, but not when CMG unloading was inhibited with p97-i (Figure 5B, compare lanes 2 and 4). Interestingly, chromatin-bound TRAIP did not increase in the presence of B1-CDK1 and p97-i compared to the level observed before B1-CDK1 addition (Figure 5B, compare lanes 1 and 4). The data suggest that TRAIP is normally part of the replisome and that it is activated by mitotic CDK, whereupon it promotes MCM7 ubiquitylation and CMG unloading.

### TRAIP promotes fork progression through a LacR array

As discussed above, p97-i not only prevented the collapse of LacR-stalled forks in mitotic extracts (Figure 4C), but also promoted the conversion of theta structures normally seen in interphase extract (Figure 4C, lanes 6–10) into mature replication products--open circular (OC) and supercoiled plasmid (SC) monomers (Figure 4C, lanes 16–20). Therefore, when CMG unloading is blocked in mitotic extracts, the replisome progresses through the LacR array more efficiently than in interphase extract. We wondered whether this enhanced fork progression depends on TRAIP. To this end, we combined p97-i treatment with TRAIP depletion. In this setting, theta structures accumulated, and the appearance of mature replication products was strongly reduced (Figure 5E, compare lanes 7–12 and 19–24), indicating that TRAIP promotes efficient replication fork progression through a LacR array. Thus, our data suggest that TRAIP-dependent CMG ubiquitylation not only promotes CMG unloading but also more efficient disruption of replication barriers when CMG unloading is blocked.

### Fork breakage in mitotic extracts is distinct from programmed incisions during ICL repair

The breakage of single stalled forks in mitotic extracts shown here is similar to the breakage of forks that have converged on cisplatin ICLs in interphase egg extracts in that both events require TRAIP-dependent CMG unloading (Wu et al., submitted). We therefore asked whether mitotic fork breakage also requires FANCI-FANCD2, XPF-ERCC1, or SLX1-SLX4, which promote DNA incisions during ICL repair. As shown in Figures S5F and S5G, immunodepletion of FANCI-FANCD2 did not prevent mitotic ARP formation on LacR plasmid, nor did depletion of SLX4, XPF, or MUS81 (data not shown). We speculate that there might be redundancy among SLX1, XPF, and MUS81 for mitotic fork breakage, or that other nucleases are involved. Our results indicate that while ICL incisions and B1-CDK1-dependent replication fork collapse both require TRAIP-dependent CMG unloading, these processes are otherwise mechanistically distinct.

### TRAIP promotes CMG unloading from terminated replisomes in mitosis

In S phase, CMG unloading during replication termination requires the E3 ubiquitin ligase CRL2^Lrr1^. However, in worms lacking CRL2^Lrr1^, CMGs persist on chromatin until late prophase, when they are unloaded from chromatin by p97 (Sonneville et al., 2017). This observation indicates that an alternative ubiquitylation pathway acts to unload terminated CMGs in mitosis, but the relevant E3 ubiquitin ligase has not been identified. To determine whether TRAIP is involved in this pathway, we first addressed whether *Xenopus* egg extracts recapitulate mitotic unloading of CMGs that have undergone replication termination. To this end, we replicated a plasmid in interphase egg extracts in the presence of Cullin inhibitor MLN-4924 (Cul-i). In this condition, DNA synthesis went to completion (Figure S6A), but CMG unloading was blocked due to inhibition of CRL2^Lrr1^ (Figure 6A, compare lanes 1 and 2; (Dewar et al., 2017)). Importantly, upon the addition of B1-CDK1, CMG was unloaded despite the presence of Cul-i (Figure 6A, lane 6), and this unloading was blocked by p97-i (Figure 6A, lane 8). Therefore, as seen in worms, frog egg extracts support CRL2^Lrr1^-independent CMG unloading in a mitotic environment. Interestingly, in the presence of p97-i, MCM7 was ubiquitylated even more extensively than in interphase extract (Figure 6A, compare lanes 7–8 and 3–4 and Figure S6B, compare lanes 5–6 and 1–2). The hyper-ubiquitylation was insensitive to Cul-i (Figure 6A, lane 8), consistent with it being CRL2^Lrr1^-independent. Importantly, TRAIP depletion inhibited B1-CDK1-induced CMG unloading from terminated forks (Figure 6B, compare lanes 1 and 4 and Figure S6C, compare lanes 1 and 4) and MCM7 hyper-ubiquitylation (Figure 6B, compare lanes 2 and 5). These defects were reversed by rTRAIP^WT^ but not rTRAIP^R18C^ (Figures 6B and S6C). Therefore, in the absence of CRL2^Lrr1^ activity, TRAIP promotes an alternative pathway to unload terminated CMGs in mitosis. Together, our results suggest that in mitosis, TRAIP removes all CMGs from chromatin, whether they have terminated or stalled (Figure S6D), with the latter case leading to fork breakage and complex end joining events (Figure 7).

**Figure 6.**
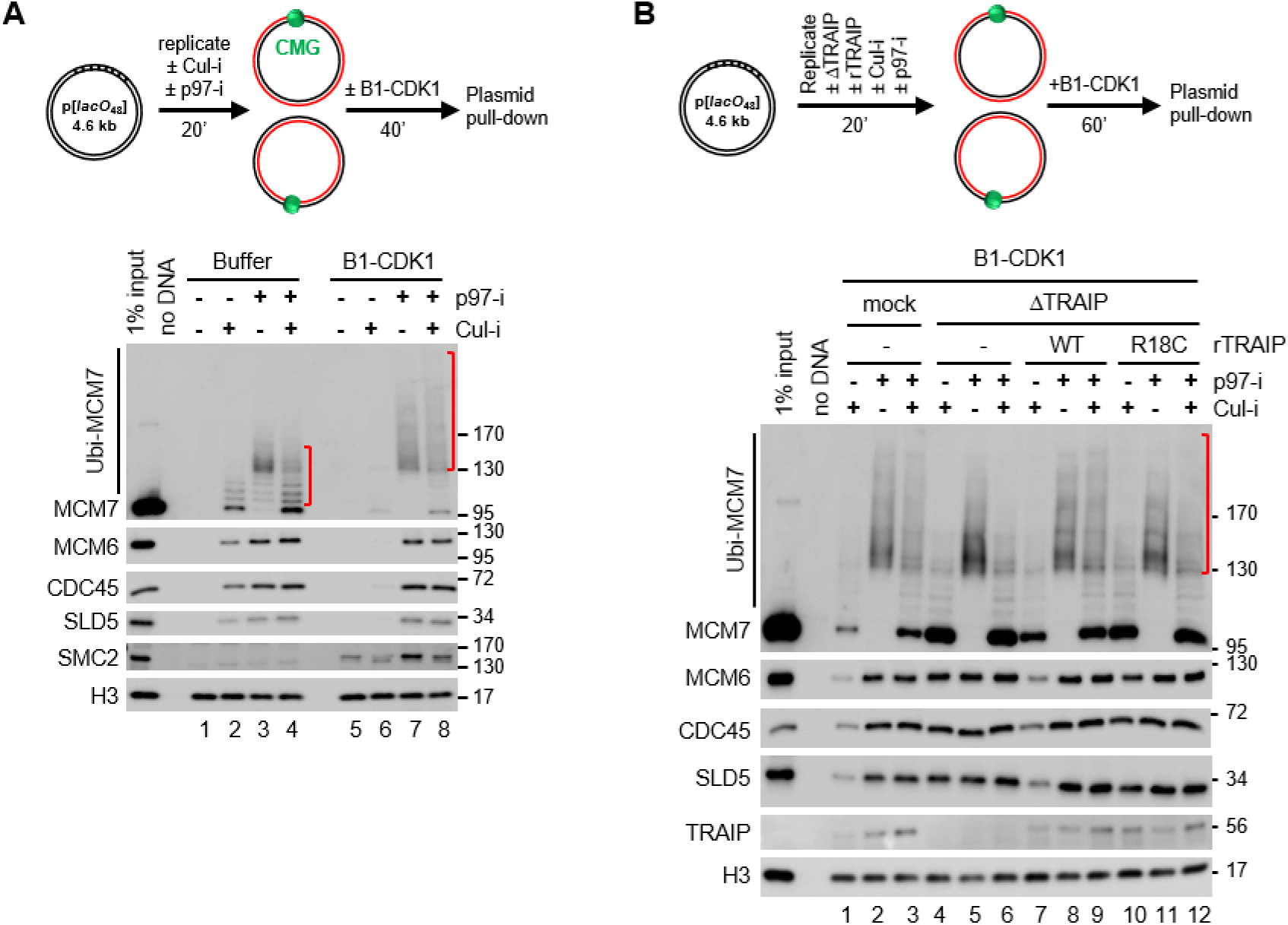
TRAIP mediates unloading of terminated CMGs in mitosis. **(A)** p[*lacO***_48_**] plasmid, in the absence of LacR, was replicated and treated as schemed. Chromatin-bound proteins were recovered and blotted with the indicated antibodies. Red brackets indicate the levels of MCM7 ubiquitylation. **(B)** p[*lacO***_48_**] plasmid, in the absence of LacR, was replicated in mock-depleted or TRAIP-depleted egg extracts supplemented with or without rTRAIP**^WT^** (~4-fold of endogenous TRAIP), or rTRAIP**^R18C^** (~9-fold of endogenous TRAIP), followed by indicated treatments. Chromatin-bound proteins were recovered and blotted with the indicated antibodies. Red bracket indicate the level of MCM7 ubiquitylation. See also **Figure S6**.

**Figure 7.**
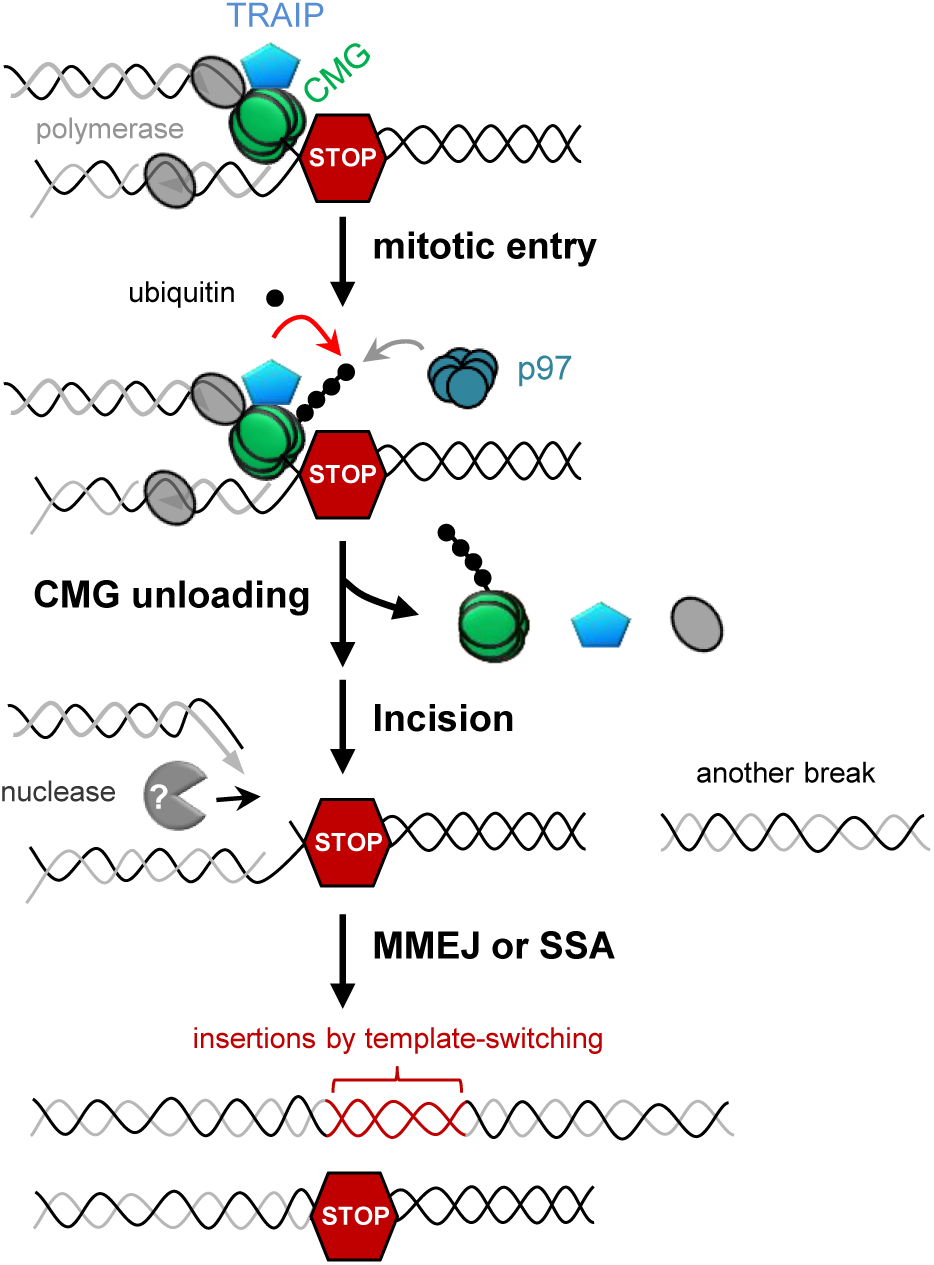
Model of CMG unloading, fork breakage and complex DNA rearrangements upon premature mitotic entry. When a replication fork encounters a replication barrier (indicated as a red hexagonal STOP sign), the replisome containing CMG and TRAIP is stably stalled during interphase. Upon mitotic entry, E3 ubiquitin ligase TRAIP is activated (directly or indirectly) to cause CMG ubiquitylation on MCM7 subunit, which in turn triggers CMG extraction/unloading from chromatin by CDC48/p97 ATPase. Loss of CMG leads to incision by so far unknown DNA nuclease(s), followed by error-prone DSB repair by MMEJ and/or SSA, which results in complex DNA rearrangements such as deletions and insertions from template-switching events.

## DISCUSSION

Numerous lines of evidence indicate that when cells enter mitosis before DNA replication is complete, replication forks collapse and break. However, the mechanism of collapse and how it affects genome stability remain obscure. Our data provide direct evidence that TRAIP-dependent replisome disassembly causes fork breakage, and they suggest a new model for the avoidance of genome instability when cells enter mitosis with unreplicated DNA.

Our findings suggest that TRAIP promotes CMG unloading in diverse contexts. In mitosis, TRAIP targets both stalled CMGs, which encircle ssDNA, and terminated CMGs, which probably encircle dsDNA (Figure S6Di and ii; (Dewar et al., 2015)). We propose that in the presence of mitotic CDK, TRAIP promotes the ubiquitylation and unloading of all CMGs, regardless of their configuration on DNA. Interestingly, TRAIP also functions in interphase: when two forks converge on an ICL, TRAIP is required to ubiquitylate CMG, and ubiquitylated CMG in turn dictates the choice between two mechanisms of ICL repair (Wu et al., submitted; Figure S6Diii). Although TRAIP promotes CMG ubiquitylation in multiple contexts, it does not target CMGs that have terminated in interphase, a function performed by CRL2^Lrr1^ (Figure S6Div; Dewar et al. 2017), nor does it appear to target CMG at single moving or stalled forks in interphase, which would cause premature fork collapse. Thus, TRAIP appears to be more selective in interphase than in mitosis, and future work will explore the basis of this difference.

There is currently no consensus on how replisome disassembly relates to replication fork collapse (Cortez, 2015; Toledo et al., 2017). *A priori*, the simplest mechanism of fork collapse would involve the loss of an essential replisome component that cannot be reloaded in S phase. If such a component also protects the fork, its loss would inevitably also cause breakage. MCM2–7 is the prime candidate, as it is the only known replication factor that cannot be loaded *de novo* in S phase (Deegan and Diffley, 2016). Consistent with this idea, we find that unloading of CMG (whose core component is MCM2–7) precedes fork breakage, and inhibition of CMG unloading via TRAIP depletion or p97-i addition suppresses breakage. To rigorously determine whether MCM7 ubiquitylation is necessary to promote CMG unloading and breakage, it will be important to mutate relevant ubiquitylation sites in MCM7. However, our mass spectrometry analysis has not identified ubiquitylated lysine residues in MCM7 (data not shown). Nevertheless, we provide the first evidence that CMG unloading is causally linked to replication fork breakage. We propose that loss of CMG might represent a common trigger of fork collapse that also leads to breakage due to exposure of the fork to one or more nucleases. It will be interesting to determine how this pathway relates to the loss of RPA at the fork, which has also been proposed to trigger fork collapse and breakage (Toledo et al., 2013).

After stressed forks undergo breakage in mitotic extract, the newly formed DNA ends undergo two classes of joining events, as revealed by DNA sequencing. The first class involves deletions of blocks of four *lacO* sites, the repeating unit within the *lacO* array. These products are most readily explained by single-strand annealing, and they are probably favored by the highly repetitive nature of the *lacO* array. SSA is usually RAD52 dependent (Bhargava et al., 2016), and RAD52 has recently been shown to mediate DNA repair synthesis during mitosis (Bhowmick et al., 2016). However, we have not been able to test the involvement of RAD52 due to an inability to raise useful antibodies against *Xenopus* RAD52. The second class of end joining products involves multiple template-switching events that are mediated by short stretches of micro-homology, indicative of DNA Polθ-mediated DNA end joining (TMEJ (Wyatt et al., 2016)). Consistent with this idea, mitotic aberrant replication products were reduced in Polθ-depleted extracts but not when homologous recombination (HR) or classical non-homologous end joining (NHEJ) was inhibited. Our observation that broken forks appear to be processed primarily by SSA and TMEJ is consistent with the finding that HR and NHEJ are inhibited in mitosis (Figure S3A and (Hustedt and Durocher, 2016; Ochs et al., 2016; Peterson et al., 2011)). Notably, we detected only short-tract template switches typical of TMEJ. If, before end joining, template-switching events mediated by Polθ or other factors were followed by more processive DNA synthesis that is templated near the break, duplications could result that resemble copy number alterations observed in human cancer and congenital disease (Carvalho and Lupski, 2016; Leibowitz et al., 2015).

When converging forks are unable to complete DNA replication by anaphase, as seen at common fragile sites (CFS), chromosome non-disjunction and aneuploidy can result. We identify two mechanisms by which TRAIP might help avoid this outcome. First TRAIP enhances CMG’s ability to overcome replisome barriers (Figure 5E), promoting the completion of replication before anaphase. Second, if the obstacle cannot be overcome, the activation of TRAIP stimulates CMG unloading and fork breakage. We propose that breakage occurs preferentially on the two leading strand templates because these are normally protected by CMG (Fu et al., 2011) and therefore exposed after CMG dissociation (Figure S7). In this scenario, one intact daughter chromosome is immediately restored by gap filling, and the other is regenerated via joining of the two broken ends, albeit with sister chromatid exchange and at the cost of of a small deletion (Figure S7, left branch). Importantly, this mechanism avoids the formation of acentric and dicentric chromosomes that would result from random breakage of the forks (Figure S7, right branch) and thus helps account for the fact that breakage at CFS is mostly beneficial (Bhowmick and Hickson, 2017; Minocherhomji et al., 2015; Naim et al., 2013; Ying et al., 2013). Strikingly, CFS expression induces chromosomal alterations that exhibit key features expected of our model, including submicroscopic deletions covering the CFS locus, microhomologies at the breakpoint junctions, and a very high frequency of sister chromatid exchanges (Glover et al., 2017) (Figure S7, left branch). Unlike our biased breakage and end joining model, break-induced replication models of CFS expression (Bhowmick et al., 2016; Minocherhomji et al., 2015) do not readily account for the high incidence of sister chromatid exchanges at CFS, and they would not be beneficial at CFS located distant from chromosome ends.

We speculate that TRAIP-dependent CMG unloading contributes to other genome instability phenomena that were previously linked to mitotic DNA replication. These include: chromosome breakage that occurs when cells in the S and M phases are fused (Duelli et al., 2007; Johnson and Rao, 1970; Rao et al., 1982), or when mitotic CDK is prematurely activated in S phase by WEE1 inhibition (Dominguez-Kelly et al., 2011; Duda et al., 2016); and chromothripsis in micronuclei that are still engaged in replication when they enter mitosis (Crasta et al., 2012; Leibowitz et al., 2015; Ly et al., 2017). In these cases, massive chromosomal breakage is deleterious as it leads to genome instability or cell death. Notably, chromosome fragmentation in the presence of WEE1 inhibitor and common fragile site expression are both MUS81-dependent (Dominguez-Kelly et al., 2011; Duda et al., 2016; Naim et al., 2013; Ying et al., 2013). In contrast, fork breakage in our experiments was not inhibited by MUS81 depletion. Whether this reflects a real difference in these processes, incomplete MUS81 depletion in extracts, or greater redundancy with other nucleases in extracts remains to be determined. It will be interesting to test the hypothesis that TRAIP underlies many different genome instability phenomena in mitosis.

We showed that in the absence of CRL2^Lrr1^ activity, TRAIP triggers the unloading of terminated CMGs in mitosis. Therefore, TRAIP likely represents the activity that removes CMGs from late prophase chromosomes in LRR-1-deficient worms (Sonneville et al., 2017). Terminated CMGs that remain on chromatin probably encircle dsDNA (Dewar et al., 2015) and thus may prevent strand separation during transcription or replication in the next cell cycle. Thus, we propose that TRAIP-dependent unloading of terminated CMGs that evaded the action of CRL2^Lrr1^ may also promote genome maintenance. Whether the dwarfism phenotype observed in patients with hypomorphic TRAIP mutations results from defective ICL repair (Wu et al., submitted), defective CMG unloading from stalled forks in mitosis, persistence of a few terminated CMGs into the next cell cycle, or the absence of other TRAIP-dependent processes remains to be established.

Although it had been widely thought that the checkpoint kinase ATR supports cell viability and suppression of replication fork collapse via phosphorylation of proteins at the fork, no ATR substrates have been identified that definitively validate this mechanism (Cortez, 2015; Pasero and Vindigni, 2017; Saldivar et al., 2017). An alternative view is that the primary role of ATR in stabilizing forks is indirect (Toledo et al., 2017). Thus, it has been proposed that ATR inhibition of late origin firing prevents exhaustion of the nuclear RPA pool, causing fork deprotection and breakage (Toledo et al., 2013). Another idea is that suppression of mitotic entry is the means by which ATR stabilizes forks, including in the absence of exogenous replication stress (Eykelenboom et al., 2013; Ragland et al., 2013; Ruiz et al., 2016). Consistent with the latter model, ATR is not required to stabilize stalled DNA replication forks in egg extracts that are permanently arrested in interphase (Luciani et al., 2004). Moreover, we show that when stressed forks are exposed to mitotic CDK, forks break, even without ATR inhibition or RPA depletion. Collectively, our data are most consistent with the idea that there is no intrinsic requirement for ATR in stabilizing forks, as long as these are not exposed to mitotic CDK activity. It will be interesting to determine whether mitotic entry and RPA exhaustion activate distinct programs of replication fork collapse and breakage.

In summary, our data suggest that when TRAIP becomes active in mitosis, a short temporal window opens in which replication forks can overcome remaining obstacles and terminate. The window closes when all CMGs are ubiquitylated and extracted from chromatin. Normally, CMG removal and fork breakage promotes chromosome segregation and genome integrity, but when too many forks are present, massive DNA fragmentation results, leading to cell death or transformation. Collectively, our results suggest that TRAIP serves a crucial role in minimizing the conflict between incomplete DNA replication and mitosis.

## ACKNOWLEDGMENTS

We thank James Dewar, Emily Low, Justin Sparks, Kyle Vrtis, Daniel Finley, Puck Knipscheer, and Jan-Michael Peters for experimental protocols or reagents. We thank Alan D’Andrea, Randy King, Ralph Scully, Karim Labib, and members of the Pellman and Walter laboratories for helpful discussion and critical reading of the manuscript. R.A.W. was supported by postdoctoral fellowship 131415-PF-17-168-01-DMC from the American Cancer Society. D.P. was supported by NIH grant CA213404. J.C.W. was supported by NIH grants GM080676 and HL098316. D.P. and J.C.W. are investigators of the Howard Hughes Medical Institute.

## AUTHOR CONTRIBUTIONS

D.P. initiated the project. L.D., D.P., and J.C.W. designed the experiments, interpreted the results, and prepared the manuscript. O.V.K. contributed Figures 6 and S6A-C; R.A.W. contributed rTRAIP^WT^ and rTRAIP^R18C^ proteins; L.D. performed all other experiments.

## DECLARATION OF INTERESTS

The authors declare no competing interests.

## METHODS

No statistical methods were used to predetermine sample size. All experiments were performed at least twice independently using separate preparations of *Xenopus* egg extracts. A representative result is shown.

### Protein purification

To purify biotinylated LacR, the LacR-Avi expressing plasmid pET11a[LacR-Avi] (Avidity, Denver, CO) and biotin ligase expressing plasmid pBirAcm (Avidity, Denver, CO) were co-transformed into T7 Express cells (New England Biolabs). Cultures were supplemented with 50 mM biotin (Research Organics, Cleveland, OH). Expression of LacR-Avi and the biotin ligase was induced by addition of IPTG (Isopropyl β-D-thiogalactoside, Sigma, St. Louis, MO) to a final concentration of 1 mM. Biotinylated LacR-Avi was then purified as described (Dewar et al., 2015). BRC (a ~35 amino acid peptide derived from BRCA2 that binds RAD51) and BRC^***^ (BRC peptide with mutations at RAD51 binding sites), a gift of K. Vrtis, were purified as reported (Long et al., 2011). rTRAIP and rTRAIP-R18C were expressed from a 6xHis-SUMO plasmid in bacteria and purified as described (Wu et al. submitted). Other proteins used in this study were Cyclin B1-CDK1 (Life Technologies Cat #PR4768C and EMD Millipore Cat #14–450M) and Cyclin E-CDK2 (EMD Millipore Cat #14–475). USP21 was a gift from D. Finley.

### DNA constructs

The 4.6 kb p[*lacO***_48_**] plasmid (a generous gift of K. Vrtis) contains an array of 48 *lacO* sites which can be bound by the *lac* repressor (LacR) to form replication barriers. The pDPC plasmid (4.3 kb), a generous gift of J. Sparks, was constructed based on a previous protocol (Duxin et al., 2014). Control plasmid (pControl) used in Figure 1G has the same DNA sequence as pDPC, but lacks crosslinks.

### *Xenopus* egg extracts and DNA replication

Egg extracts were prepared using *Xenopus laevis* (Nasco Cat #LM0053MX). All experiments involving animals were approved by the Harvard Medical School Institutional Animal Care and use Committee (IACUC) and conform to relevant regulatory standards. *Xenopus* egg extracts including Low Speed Supernatant (LSS), High Speed Supernatant (HSS), and Nucleoplasmic egg extract (NPE) were prepared as described (Blow and Laskey, 1986; Lebofsky et al., 2009).

To assess the effects of mitotic cyclins, demembranated sperm chromatin from *Xenopus laevis* males was incubated in LSS (4,000 sperms/μL LSS) for 40 minutes at room temperature to form nuclei. The reactions were subsequently incubated with a range of concentrations of mitotic B1-CDK1. Nuclear envelope integrity and chromatin condensation were monitored by microscopy after Hoechst staining (see below). The concentration (50 ng/μL) that triggered nuclear envelopment breakdown and chromosome condensation was chosen to trigger mitotic entry in subsequent experiments.

For interphase DNA replication, sperm chromatin or plasmid DNA was first incubated in HSS (final concentration of 7.5–15.0 ng DNA/μL HSS) for 30 minutes at room temperature to license the DNA for replication (“licensing”), followed by the addition of 2 volumes of NPE to initiate CDK2-dependent replication. To radiolabel the nascent strands during replication, NPE was supplemented with trace amounts of [α-^32^P]-dATP. Mitotic DNA replication was performed essentially as described (Prokhorova et al., 2003). Briefly, after 30 minutes, 0.9 volumes of licensing reaction was incubated with 0.1 volumes of mitotic B1-CDK1 for 30 minutes at room temperature, followed by addition of 2 volumes of NPE. In the “licensing” mixture, the concentration of B1-CDK1 was 50 ng/μL, and its concentration in the final replication reaction was 16.7 ng/μL. Unless stated otherwise, the ‘0 minute’ time point refers to the moment of NPE addition. 2 μL aliquots of replication reaction were stopped with 5 μl of stop solution A (5% SDS, 80 mM Tris pH8.0, 0.13% phosphoric acid, 10% Ficoll) supplemented with 1 μl 20 mg/ml Proteinase K (Roche, Nutley, NJ). Samples were incubated for 1 hour at 37°C prior to electrophoresis on a 0.9% native agarose gel. Gels were dried and radioactivity was detected using a phosphorimager (Lebofsky et al., 2009).

To induce replication fork stalling using LacR, one volume of p[*lacO***_48_**] (200 ng/μL) was incubated with one volume of recombinant LacR (36 μM) for 45–60 minutes at room temperature. Next, 0.1 volumes of the mixture was combined with 0.9 volumes of HSS for licensing, followed by addition of 2 volumes of NPE for initiation of replication. To inhibit the binding of LacR to the *lacO* array, IPTG was added to NPE to a final concentration of 10 mM and incubated for 15 minutes prior to use in replication (Figure 1E) or added into replication reactions after fork stalling (Figure 4G) at the indicated time.

For replication assays with inhibitors, NPE was supplemented with inhibitors for 15 minutes at room temperature before addition to the licensing mixture. Inhibitors were used at the following final concentrations in replication reaction: Aphidicolin (Sigma Cat #A0781–5MG), 2.2 μM or 0.97 μM, as indicated; CDC7 inhibitor PHA-767491 (Sigma Cat #PZ0178), 266 μM; p97 inhibitor NMS-873 (Sigma Cat #SML1128–5MG), 266 μM; DNA-PKcs inhibitor NU-7441, 133 μM; BRC or BRC^***^, 1 μg/μL; Cullin inhibitor MLN-4924 (Active Biochem Cat #A-1139), 266 μM. For the Cdk1 inhibition assay in Figure S2B, CDK1 inhibitor RO-3306 (EMD Millipore Cat #217699–5MG) was incubated with the replication reaction containing stalled replication forks for 5 minutes before the addition of B1-CDK1.

### Immunodepletion and Western blotting

Immunodepletions using antibodies against *Xenopus laevis* FANCD2 (Knipscheer et al., 2009), FANCI (Duxin et al., 2014), SMC2 (antigen: Ac-CSKTKERRNRMEVDK-OH, New England Peptide), TRAIP (antigen: Ac-CTSSLANQPRLEDFLK-OH, New England Peptide), Polθ (antigen: residues 1212 to 1506, Abgent), and RAD51 (Long et al., 2011) were performed as described previously (Budzowska et al., 2015). Briefly, Protein A Sepharose Fast Flow beads (GE Healthcare) were incubated with antibodies at 4°C overnight. For mock depletion, an equivalent quantity of nonspecific rabbit IgGs was used. Five volumes of pre-cleared HSS or NPE were then mixed with one volume of the antibody-bound sepharose beads. For FANCI-D2 depletion of HSS and NPE, two rounds of depletion using both FANCI and FANCD2 antibodies were performed at room temperature for 20 minutes each. Depletions for other proteins were performed at 4°C, with two rounds for HSS and three rounds for NPE. For each round, a mixture of antibody-bound beads and egg extract was rotated on a wheel for 40 minutes. Immunodepleted extracts were collected and used immediately for DNA replication. Depletion efficiency was assessed by Western blotting. Western blots from depletion or plasmid/sperm chromatin pull-downs were probed using antibodies against SMC2, TRAIP, FANCI (Duxin et al., 2014), FANCD2 (Knipscheer et al., 2009), MCM7 (Dewar et al., 2017), MCM6 (Dewar et al., 2017), RAD51 (Long et al., 2011), ORC2 (Dewar et al., 2017), CDC45 (Walter and Newport, 2000), SLD5 (Dewar et al., 2017) and Histone H3 (Cell Signaling Technology #9715S).

### Sperm chromatin spin-down assay

Sperm chromatin spin-down was performed as previously described (Raschle et al., 2015). Briefly, chromatin and associated proteins were isolated by centrifugation through a sucrose cushion, washed three times, resuspended in 2x SDS sample buffer (100 mM Tris pH 6.8, 4% SDS, 0.2% bromophenol blue, 20% glycerol, 10% β-mercaptoethanol) and boiled at 95°C for 3–5 minutes. In Figure S3A, chromatin was spun down 20 minutes after NPE addition for the Buffer control and at 9 minutes after NPE addition for the B1-CDK1 treatment (final concentration, 16.7 ng/μL), at which point replication was ~50% complete for both reactions. In Figure S1D, chromatin and associated proteins were isolated from HSS.

### Plasmid pull-down assay

Plasmid pull-down assays were performed as described (Budzowska et al., 2015). Briefly, streptavidin-coupled magnetic beads (Dynabeads M-280, Invitrogen; 6 μl beads slurry per pull-down) were washed three times with wash buffer 1 (50 mM Tris pH 7.5, 150 mM NaCl, 1 mM EDTA pH 8, 0.02% Tween-20). Biotinylated LacR was incubated with the beads (12 pmol per 6 μL beads) at room temperature for 40 min. The beads were then washed four times with pull-down buffer 1 (10 mM Hepes pH 7.7, 50 mM KCl, 2.5 mM MgCl_2_, 250 mM sucrose, 0.25 mg/mL BSA, 0.02% Tween-20) and resuspended in 40 μL of the same buffer. At the indicated times, 4 μL samples of the replication reaction were withdrawn and gently mixed with Biotin-LacR-coated beads. The suspension was immediately placed on a rotating wheel and incubated for 30–60 minutes at 4°C. The beads were washed three times with wash buffer 2 (10 mM Hepes pH 7.7, 50 mM KCl, 2.5 mM MgCl_2_, 0.25 mg/mL BSA, 0.03% Tween-20). The beads were resuspended in 40 μL of 2× SDS sample buffer and boiled at 95°C for 3–5 minutes. Chromatin-bound proteins were separated by SDS-PAGE and analyzed by Western blotting.

### De-ubiquitination assay

Plasmid pull-downs were performed as described above, except that after the wash steps with wash buffer 2, chromatin-bound proteins were resuspended in 20 μL of USP21 buffer (150 mM NaCl, 10 mM DTT, 50mM Tris pH 7.5) and split into two 10 μL aliquots. Each aliquot was incubated with 1 μL of the non-specific deubiquitinase USP21 or buffer at 37°C for 60 minutes. The reactions were stopped by addition of 2x SDS sample buffer and boiled at 95°C for 3–5 minutes.

### Restriction digestion

2 μL aliquots of replication reactions were stopped in 20 μL of stop solution B (50 mM Tris pH 7.5, 0.5% SDS, 25 mM EDTA), and replication products were purified as previously described (Raschle et al., 2008). Purified products were digested with restriction enzymes as *per* the manufacturer’s instructions. Digestion reactions were stopped in 0.5 volumes of stop solution C (5% SDS, 4 mg/mL Proteinase K) and incubated for 60 minutes at room temperature. Digested products were separated on a 1% native agarose gel and visualized by autoradiography.

### Sequencing

LacR-bound p[*lacO***_48_**] plasmid was replicated in the presence of mitotic B1-CDK1 for 120 minutes. Replication products were purified and digested with AlwNI (single cut on the parental DNA) for 60 minutes at 37°C, as described above. After separation on a 0.9% native agarose gel, bands smaller than the 4.6 kb full-length linear fragment were extracted and self-ligated with T4 DNA ligase. The ligation products were transformed into *E.coli* DH5α. As a control, p[*lacO***_48_**] was replicated without LacR for 120 minutes in the presence of B1-CDK1. Replication products (containing only open circular and supercoiled species) were processed as above, and the only band (4.6 kb) after AlwNI restriction was purified for cloning. Clones from both treatments were sequenced by Sanger method with Forward primer: 5’-AAGGCGATTAAGTTGGGTAA-3’ and Reverse primer: 5’-CATGTTCTTTCCTGCGTTATCCCCTGA-3’.

### Microscopy

1 μL of nuclear assembly reactions containing LSS egg extract and sperm chromatin was mixed with 1 μL of Hoechst 3300 (2.5 μg/mL) for 5 minutes before imaging. Images in Figures S1A and S1C were single focal planes acquired by a wide field Nikon Eclipse E600 microscope equipped with a Nikon 40x Plan Apo NA 1.0 oil objective. Images in Figure S4B were maximum projections from stacks of z-series acquired with a 0.5 μm step size. Images were collected using a 60x Plan Apo NA 1.4 oil objective with a CoolSnapHQ2 CCD camera (Photometrics) on a Yokogawa CSU-22 spinning disk confocal system (Nikon Instruments, Melville, NY). Fluorophores were excited by a 405 nm laser.

### Data quantification

Autoradiographs and Western blots were quantified using ImageJ 1.48v (National Institute of Health). The quantification methods for individual results are described in the figure legends.

## SUPPLEMENTAL FIGURE LEGENDS

**Figure S1, related to Figure 1.**
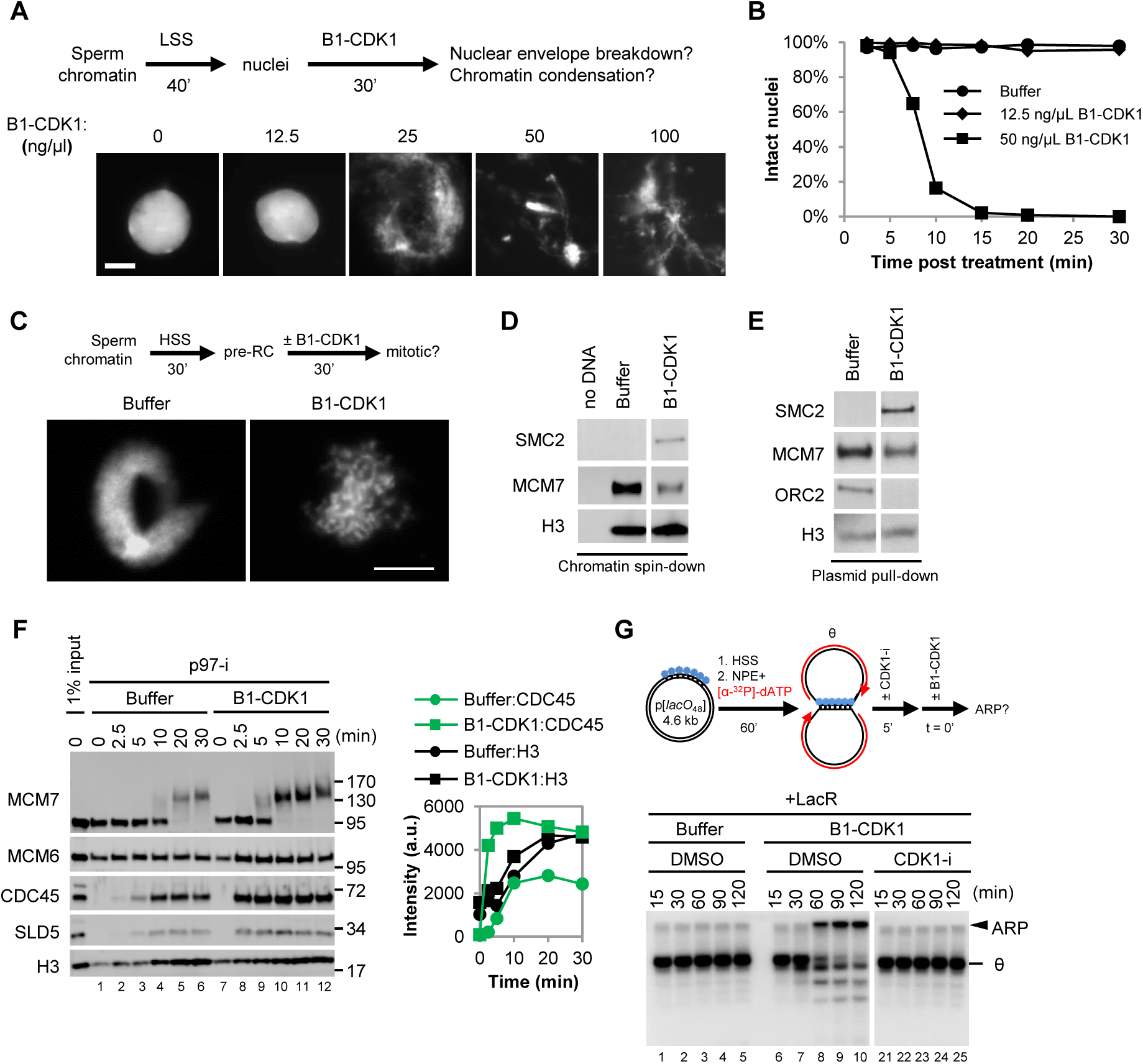
**(A)** To determine the concentration of mitotic B1-CDK1 that efficiently induces nuclear envelope breakdown and chromatin condensation, de-membranated *Xenopus* sperm chromatin was incubated in LSS (low speed supernatant) for 40 minutes to allow the formation of pseudo nuclei. The indicated final concentrations of B1-CDK1 were then added into the reactions for 30 minutes before Hoechst staining and imaging. 50 ng/μL of B1-CDK1 was sufficient to induce nuclear envelope breakdown and chromatin condensation and it was used for subsequent experiments unless otherwise indicated. Scale bar, 10 μm. **(B)** Percentage of intact nuclei remaining at the indicated time points after treatment with the indicated concentration of B1-CDK1 (n>1,000). The ‘0 minute’ time point refers to Buffer or B1-CDK1 addition. The value at each time point was normalized to the value at 0 minute in each treatment. **(C)** Chromatin condensation assay in membrane-free HSS. Sperm chromatin was incubated in HSS for 30 minutes, and then treated with 50 ng/μL of B1-CDK1 for 30 minutes followed by Hoechst staining and imaging. Scale bar, 10 μm. **(D)** Sperm chromatin spin-down assays in HSS. Sperm chromatin was incubated with HSS for 30 minutes and treated with Buffer or 50 ng/μL of B1-CDK1 for another 30 minutes. Chromatin DNA was recovered and chromatin-bound proteins were blotted with indicated antibodies. Unrelated lanes were cropped as indicated by the gap. **(E)** Plasmid pull-down assays in HSS. pBlueScript (3 kb) was incubated with HSS at a concentration of 7.5 ng/μL for 30 minutes and treated with Buffer or 50 ng/μL of B1-CDK1 for another 30 minutes. Plasmid was recovered and chromatin-bound proteins were blotted with indicated antibodies. Unrelated lanes were cropped as indicated by the gap. **(F)** Plasmid pull-down assay to assess origin firing. pBlueScript was incubated with HSS for 30 minutes and treated with buffer or 50 ng/μL of B1-CDK1 for another 30 minutes before addition of NPE. The p97 inhibitor NMS-873 (p97-i) was added into NPE (final concentration, 266 μM) and incubated for 15 minutes. Treatment of p97-i blocked the unloading of CMG helicases from chromatin and trapped ubiquitylated MCM7 on chromatin, seen as a smear. Right panel shows the quantification of the CDC45 and Histone H3 signals. Increased CDC45 loading with B1-CDK1 treatment suggested more origin firing. **(G)** LacR-bound p[*lacO***_48_**] was replicated in interphase egg extracts for 60 minutes and then treated with DMSO or Cdk1 kinase inhibitor (CDK1-i, 333 μM RO-3306) for 5 minutes before the addition of Buffer or 50 ng/μL B1-CDK1. At the indicated times, samples were withdrawn and replication products were tracked by electrophoresis and autoradiography. ARP, aberrant replication product; θ, theta structure.

**Figure S2, related to Figure 2.**
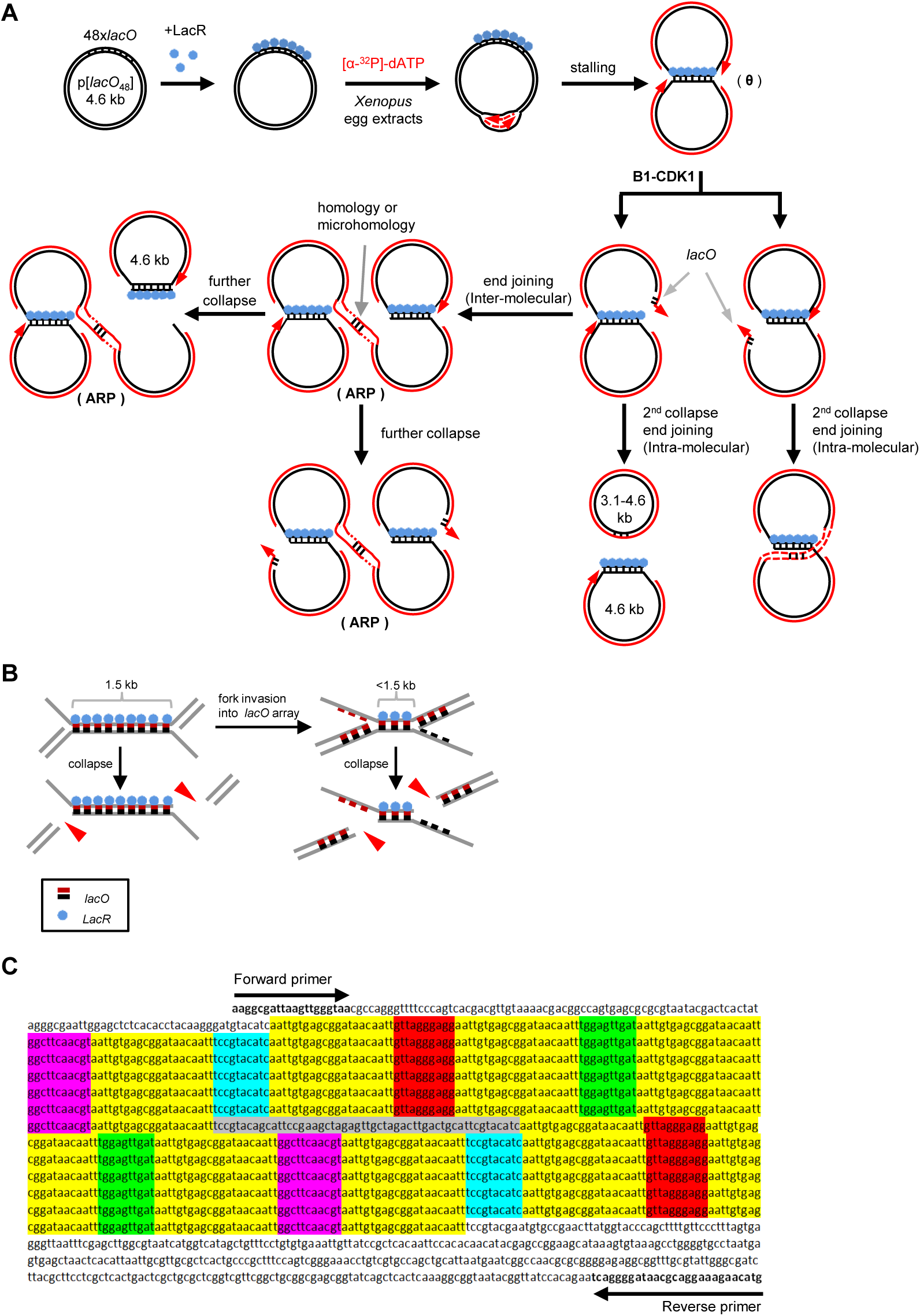
**(A)** Model for mitotic processing of replication forks stalled by *lacO*-LacR barriers, explaining the restriction analysis (Figure 2B) and sequencing data (Figure 2D). After replication fork stalling, B1-CDK1 induces fork collapse and double-strand breaks (DSBs) at the edges of the *lacO* array. The broken DNA ends, with certain number of *lacO* repeats, lead to either intra- or inter-molecular end joining. Inter-molecular end joining generates the aberrant replication products (ARPs). The initial end joining products can also be subject to cycles of fork collapse and end joining. Outcomes other than those illustrated here are possible but may not be detected because our sequencing strategy depends on the ability to recover plasmids by cloning. Although it has not been addressed whether the leading or lagging strand templates break, the results on CMG unloading described in Figure 7, as drawn (see below and text for details) favor the leading strand breakage. **(B)** Schematic of B1-CDK1-induced fork breakage at different locations in the *lacO* array. Breakage at the outer edges (left) and joining of the resulting one-ended breaks creates large deletions of the array, whereas breakage closer to the midpoint of the array causes smaller deletions (right). **(C)** Sequence and structure of the 48 *lacO* repeats in p[*lacO***_48_**]. Each *lacO* repeat is highlighted in yellow. Unique spacer sequences between *lacO* repeats are highlighted in red, green, purple and blue, respectively, as depicted in Figures 2D and 2E. The sequence in grey indicates a unique spacer in the middle of the *lacO* array. Sequencing primers used in Figure 2D are indicated.

**Figure S3, related to Figure 3.**
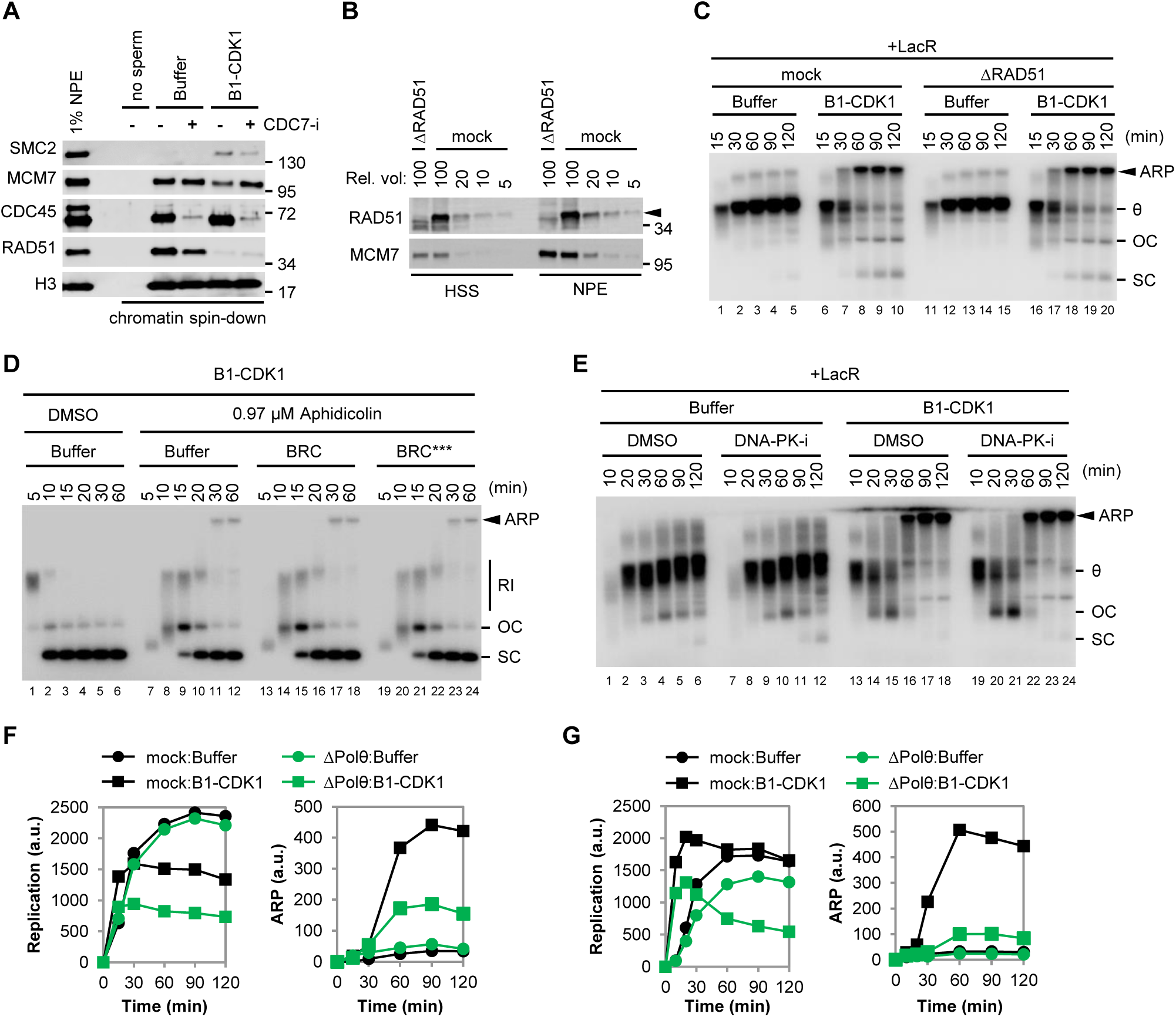
**(A)** B1-CDK1 treatment inhibits chromatin-loading of RAD51. Sperm chromatin was replicated in egg extracts and sampled when 50% replication was completed (20 minutes for Buffer and 9 minutes for B1-CDK1). To inhibit DNA replication, CDC7 inhibitor (CDC7-i, 399 μM of PHA-767491) was added to NPE and incubated for 15 minutes. Chromatin-bound proteins were recovered by chromatin spin-down and detected by blotting with indicated antibodies. **(B)** Mock-depleted and RAD51-depleted egg extracts were blotted with RAD51 and MCM7 antibodies. Serial dilutions of mock-depletion were used to assess the level of RAD51 depletion. Arrowhead indicates RAD51. **(C)** LacR-bound p[*lacO***_48_**] was replicated in mock-depleted or RAD51-depleted egg extracts in the absence or presence of B1-CDK1. **(D)** pBlueScript was replicated in egg extracts with the indicated treatments. BRC peptide binds and blocks RAD51’s interaction with BRCA2, which prevents HR-mediated DSB repair. BRC^***^ peptide harbors three mutations at RAD51 binding sites and is unable to inhibit RAD51 (Long et al., 2011). **(E)** LacR-bound p[*lacO***_48_**] was replicated with the indicated treatments. To inhibit NHEJ, a DNA-PK inhibitor (DNA-PK-i, 133 μM NU-7441) was added to NPE. **(F)** Quantification of overall DNA replication and ARP for Figure 3B. **(G)** Quantification of overall DNA replication and ARP for Figure 3C. In (C-E), ARP, aberrant replication product; θ, theta structure; OC, open circle; SC, supercoil; RI, replication intermediate.

**Figure S4, related to Figure 4.**
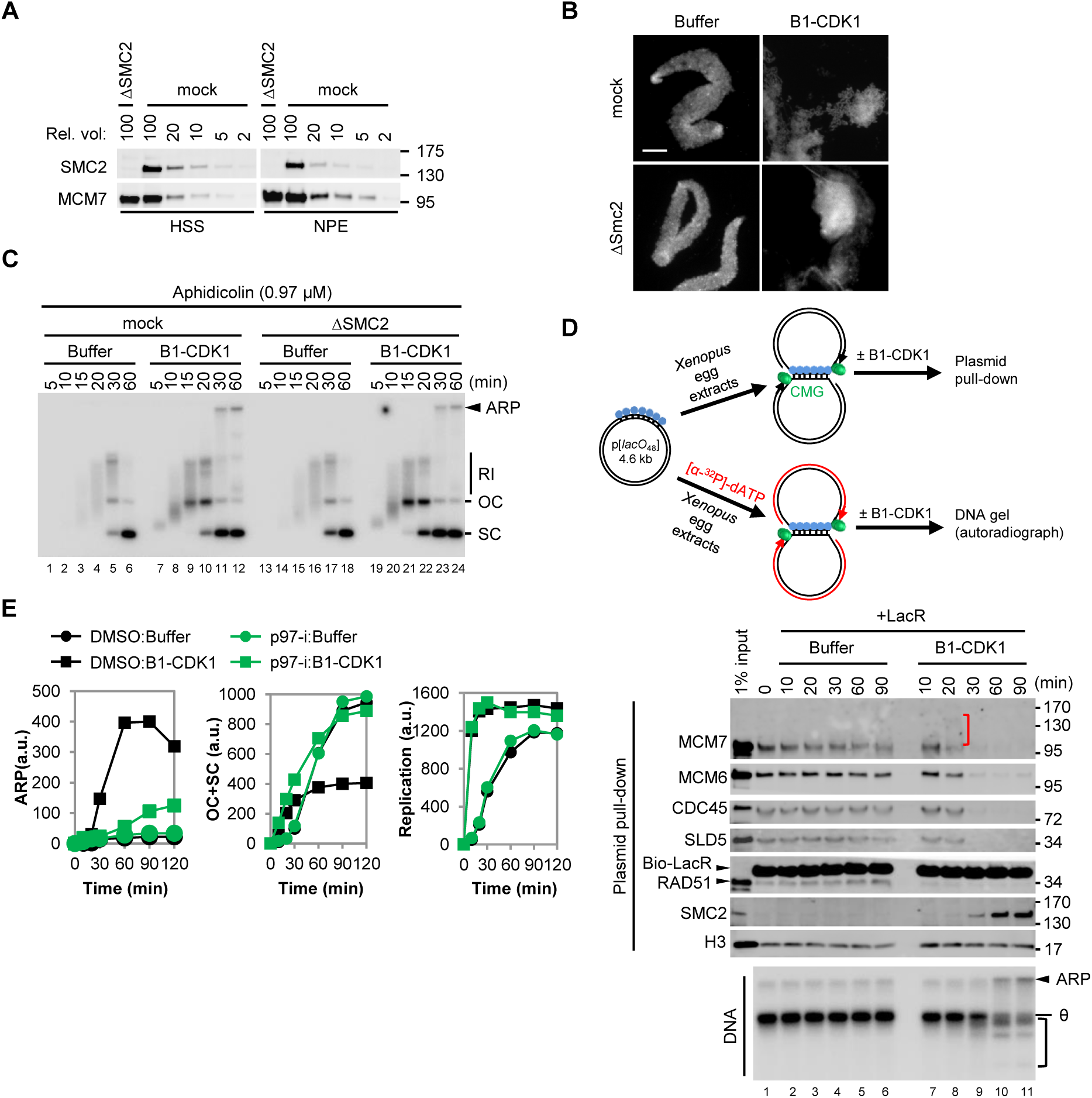
**(A)** Mock-depleted and SMC2-depleted *Xenopus* egg extracts were blotted for SMC2 and MCM7 alongside a serial dilution of mock-depleted extracts. **(B)** Effect of SMC2 depletion on B1-CDK1-induced chromatin condensation in HSS. Sperm chromatin was incubated in mock-depleted or SMC2-depleted HSS with Buffer or B1-CDK1 for 30 minutes prior to Hoechst staining and imaging. Scale bar, 10 μM. **(C)** pBlueScript was replicated in mock-depleted or SMC2-depleted egg extracts with a low dose of aphidicolin in the absence or presence of B1-CDK1. The absence of SMC2 had no effect on mitotic ARP formation. **(D)** A time course to relate the timing of CMG unloading to replication fork collapse and ARP formation during replication with B1-CDK1. LacR-bound p[*lacO***_48_**] was replicated in egg extracts for 30 minutes before the addition of Buffer or B1-CDK1. Plasmid pull-downs were performed from “cold” reactions lacking radio-labeled nucleotides in parallel with “hot” reactions containing [α-^32^P]-dATP. Plasmid pull-down samples were blotted for indicated proteins. Replication products were detected by autoradiography after gel electrophoresis. The red bracket indicates ubiquitylated MCM7, which is detectable before the appearance of the ARP. The black bracket marks potential collapsed replication forks with the B1-CDK1 treatment. **(E)** Quantification of ARP, OC+SC, and overall DNA replication during replication of pDPC in Figure 4D. In (C) and (D), RI, replication intermediate; ARP, aberrant replication product; OC, open circle; SC, supercoil; θ, theta structure.

**Figure S5, related to Figure 5.**
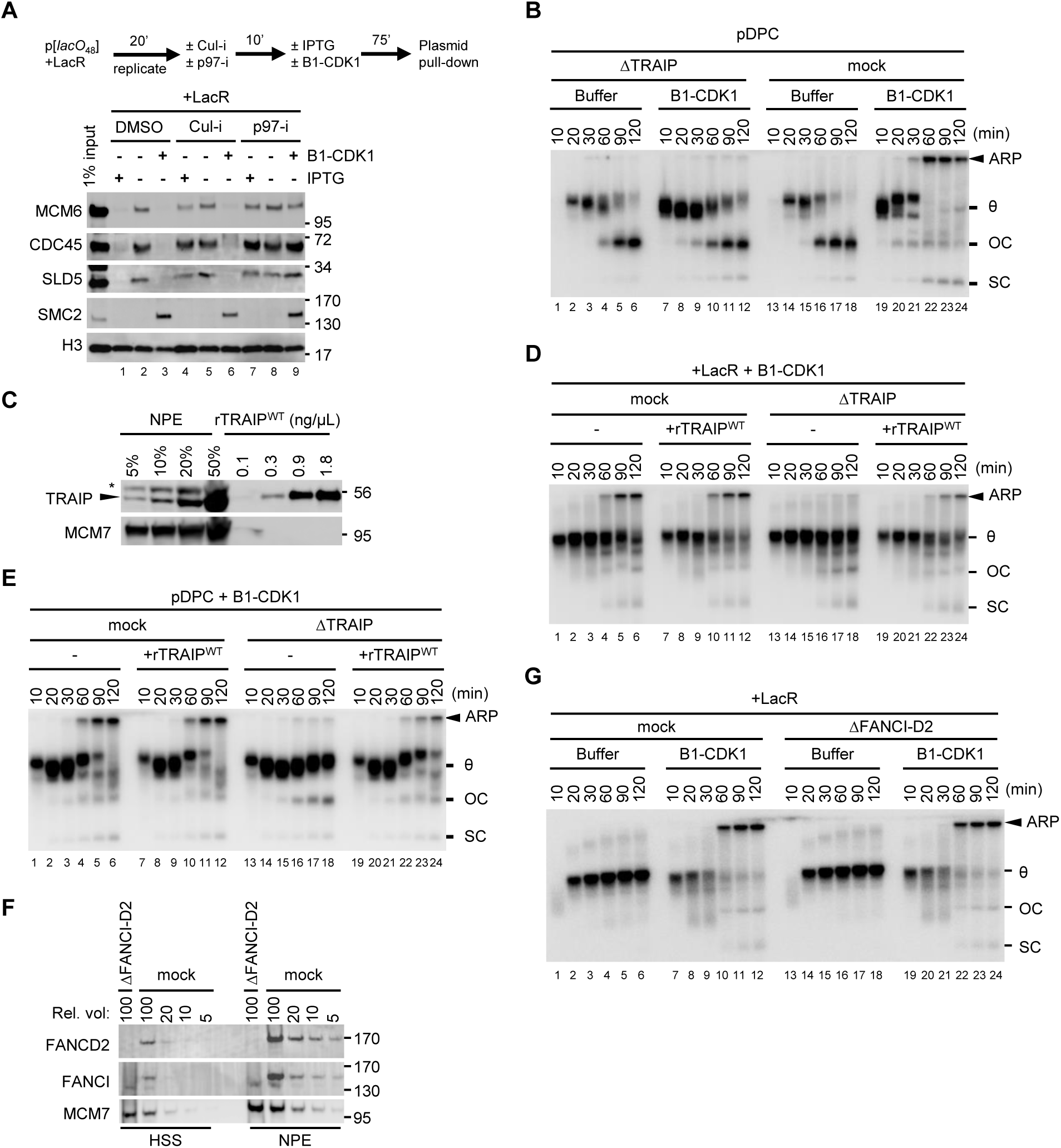
**(A)** LacR-bound p[*lacO***_48_**] was replicated and treated as schemed. Chromatin-bound proteins were recovered and blotted with the indicated antibodies. IPTG was used to release LacR from *lacO* array therefore induce replication termination. Cul-i was used to inhibit CRL2^Lrr1^-dependent CMG ubiquitylation during interphase replication termination. **(B)** pDPC was replicated in mock-depleted or TRAIP-depleted egg extracts in the presence or absence of B1-CDK1. **(C)** Serial dilutions of NPE and rTRAIP^WT^ purified from *E. coli* were blotted with TRAIP and MCM7 antibodies. Arrow head marks TRAIP signal and asterisk indicates a background band in NPE. The concentration of TRAIP in NPE is 3.0–4.5 ng/μL. **(D)** LacR-bound p[*lacO***_48_**] was replicated in mitotic mock-depleted or TRAIP-depleted egg extracts with or without rTRAIP^WT^ as indicated. rTRAIP^WT^ was added to NPE at endogenous level (3.6 ng/μL). Matched buffer was added to reactions without rTRAIP^WT^. **(E)** pDPC was replicated in mitotic mock-depleted or TRAIP-depleted egg extracts with or without rTRAIP^WT^, as indicated. rTRAIP^WT^ was added to NPE at endogenous level (3.6 ng/μL). Matched buffer was added to reactions without rTRAIP^WT^. **(F)** Mock-depleted and FANCI-D2-double depleted egg extracts were blotted with indicated antibodies. Serial dilution of mock-depleted extract was used to assess the level of FANCI-D2 depletion. **(G)** LacR-bound p[*lacO***_48_**] was replicated in mock-depleted or FANCI-D2-depleted egg extracts in the absence or presence of B1-CDK1. The depletion of FANCI-FANCD2 had no effect on ARP formation.In (B), (D), (E) and (G), ARP, aberrant replication product; θ, theta structure; OC, open circle; SC, supercoil.

**Figure S6, related to Figure 6.**
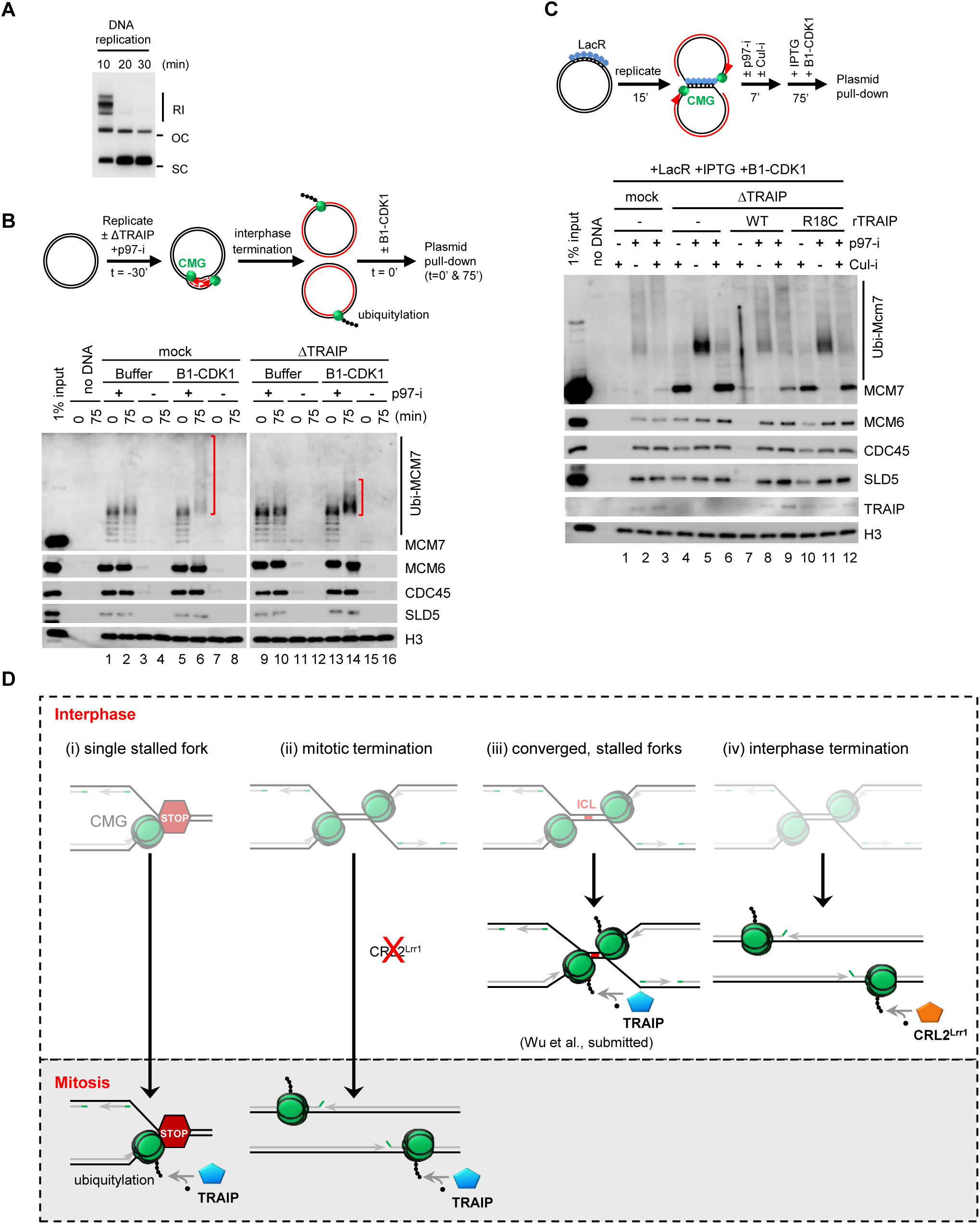
**(A)** p[*lacO***_48_**], in the absence of LacR, was replicated in egg extracts used in Figures 6A and 6B. DNA replication was complete in 20 minutes. RI, replication intermediate; OC, open circle; SC, supercoil. **(B)** A 3.1 kb plasmid (pJD152 in (Dewar et al., 2015)) was replicated in mock-depleted or TRAIP-depleted extracts in the presence or absence of p97-i (to trap terminated and ubiquitylated CMGs on chromatin) followed by Buffer or B1-CDK1 treatment. Chromatin-bound proteins were recovered and blotted with indicated antibodies. Red brackets indicate the levels of MCM7 ubiquitylation. Note the dramatic smear of MCM7 ubiquitylation in the presence of B1-CDK1 in mock (compare lanes 6 and 2) and the shrinkage with TRAIP depletion (compare lanes 14 and 6). **(C)** LacR-bound p[*lacO***_48_**] plasmid was replicated in mock-depleted or TRAIP-depleted egg extracts with or without recombinant rTRAIP^WT^ (~4-fold of endogenous TRAIP), or rTRAIP^R18C^ (~9-fold of endogenous TRAIP), and treated as schemed. Chromatin-bound proteins were recovered and blotted with the indicated antibodies. **(D)** Comparison of CMG unloading pathways. Mitotic CMG unloading at single stalled fork (i) occurs when a single stalled CMG on ssDNA enters mitosis. TRAIP is activated by mitotic CDK to trigger CMG ubiquitylation. Mitotic termination (ii) occurs when CRL2^Lrr1^ is deficient (Sonneville et al., 2017). CMGs at terminated replication forks are ubiquitylated upon mitotic entry in a TRAIP-dependent manner. During interphase ICL repair (iii) (Wu et al., submitted), when two CMGs on ssDNA converge at ICL, TRAIP is activated, independent of CDK1 activity (data not shown) and promotes CMG ubiquitylation. During replication termination in interphase (iv), two CMGs bypass each other and translocate from ssDNA to dsDNA, triggering CRL2^Lrr1^-dependent CMG ubiquitylation (Dewar et al., 2015; Dewar et al., 2017; Sonneville et al., 2017). The cartoons highlight the requirement of E3 ubiquitin ligase activity rather than physical localization for CMG ubiquitylation. In contrast to CRL2^Lrr1^ which is specifically recruited to replisome during interphase replication termination, TRAIP travels with the replisome.

**Figure S7. Related to Figure 7.**
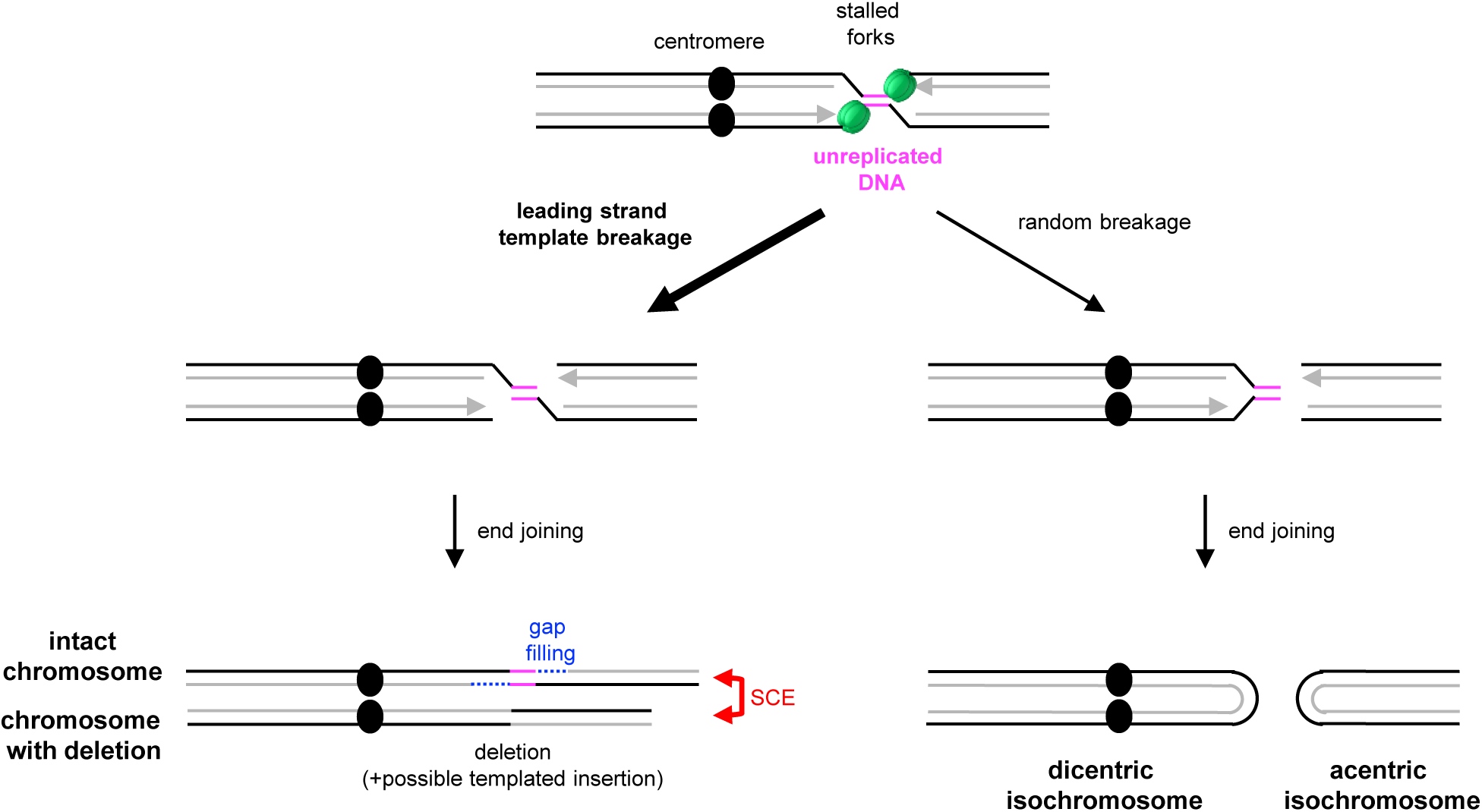
When replication forks stall on either side of a hard-to-replicate region (e.g. a common fragile site), entry into mitosis causes CMG unloading and efficient fork breakage. Because CMG binds the leading strand template, we propose that CMG unloading leads to breakage of both stalled forks on the leading strand templates (left pathway). One intact sister chromatid is rapidly restored by gap filling (dashed blue line). The other chromatid is restored by alternative end joining of the two broken ends, yielding sister chromatid exchange and a deletion that encompasses the segment of unreplicated DNA. Template switching before end joining could generate duplications at the breakpoint. In contrast, if stalled forks are broken randomly (right pathway), unproductive outcomes will be frequent, including the formation of acentric and dicentric isochromosomes (shown). Furthermore, if only one fork is broken, acentric arms can be generated (not shown).

